# A Geometrically Transient Material for Bioelectronic Implants

**DOI:** 10.1101/2025.06.17.660105

**Authors:** Selin Olenik, John D. Goodwin, Atharv Naik, Philip Coatsworth, Ioi Chit Cheung, Farbod Amirghasemi, Anies Sohi, Ehsan Abedi, Maral Mousavi, Andriy S. Kozlov, Victoria Salem, Andrew S. Cowburn, Firat Guder

## Abstract

The outstanding barrier properties of skin make it difficult to obtain reliable physiological information (especially chemical) without the use of implantable bioelectronic sensing devices to directly access the internal biology. The clinical utility of bioelectronic implants however, hinges on a key geometrical optimization problem: devices must scale-down in size to reduce surgical invasiveness while also creating enough space to integrate electronics for wireless power delivery, data exchange, and electrical/electrochemical monitoring. Here, we present a minimally invasive bioelectronic implant with a transient geometry that can be inserted subcutaneously and measures important markers, such as pH, temperature, cardiac and respiratory activity, and levels of lithium, an important element for medical applications. To produce this new class of minimally invasive and foldable implantable sensors, we developed a fabrication method that works with highly flexible substrates to enable multiple-fold miniaturization during implantation. After implantation, the implant autonomously unfolds back to its planar form for continuous wireless operation. We demonstrate the key concepts concerning implantation, operation, and removal through extensive *in vitro*, *ex vivo*, and *in vivo* animal experiments. Our approach allows for robust multiplexed monitoring using quick and suture-free insertion procedures and may provide a unique advantage in the transition towards personalized health profiles.

## Introduction

Skin is a remarkable chemical, physical, and electrical barrier. Despite being only a few millimeters thick, the outstanding barrier properties of skin make it difficult to obtain reliable physiological information from within the body non-invasively using wearable sensors, especially when trying to collect chemical information^1–7^. Non-invasive wearable sensors, therefore, exist today primarily as wellness and fitness monitors^8–10^. In contrast, high performance clinical tools such as continuous glucose or cardiac monitors prefer invasive monitoring – that is, going through the barrier of the skin – to directly access the biomarker(s) of interest and ensure precision and accuracy^11,12^.

Implantable bioelectronic sensing devices (IBSDs) have been used in niche applications of medicine for decades (for example, cochlear implants) but have had a limited range of applications. This is primarily due to the size of most implants: bulky batteries (far exceeding the size of the electronics) are required to provide long-term operation (that is, years). Larger devices require invasive surgeries for implantation which carry substantial risk to the patient, including: complications from surgery or anesthesia, increased risk of infection (and sepsis) from large incisions, and surgical re-intervention for battery replacement^13,14^. Traditional implantation procedures also require medical expertise and facilities that cannot be easily found outside major medical centers^15^. Consequently, IBSDs have been restricted to emergency and/or high-risk applications, such as the use of implantable cardiac monitors (ICMs) for detecting arrythmias^16^. To improve the clinical utility of IBSDs, therefore, it is critical to reduce the size of implants and subsequently the invasiveness of implantation to minor procedures such as injection or insertion.

The simplest method to reduce the size, hence invasiveness, of an IBSD is to eliminate the battery and power the device wirelessly. With the reduction of size, however, a critical geometric optimization problem emerges. While miniaturized 3D coil antennas have been integrated to create devices that can be inserted under the skin (< 5mm diameter), millimeter-scale antennas are inefficient and reduce the strength of power delivery and data communication^11,17^. High levels of miniaturization also negatively impact sensing: for example, the separation and size of the electrodes are critical to increasing the quality of electrical and electrochemical transduction. The size of IBSDs can also be reduced by replacing batteries with a planar antenna. Planar antennas are more popular because they provide sufficient space for the integration of off-the-shelf components in IBSDs, which accelerate development and fabrication^18–21^. The increased robustness of power delivery also allows for more complex capabilities, such as multiplexed monitoring^22,23^. To meet the requirements for operation, however, the size of the antenna – typically over 10mm in width – increases beyond the threshold for minimally invasive procedures such as insertion.

The ideal IBSD, therefore, should have a transient geometry: the implant should be as small as possible during implantation to reduce invasiveness and large when needed after implantation to allow wireless delivery of power, exchange of data, and high performance electrical/electrochemical transduction. In this regard, shape memory polymers have been explored as a substrate that can be programmed to a miniaturized form for implantation and then stimulated inside the body (by heat or light, for example) to reprogram back to its planar form^24,25^. Notably, Jiang et al. produced a wireless sensor which could be rolled to fit inside of a two-millimeter syringe for subcutaneous injection^26^. The implant lacks electronics, however, and cannot provide digital capabilities (such as data storage and processing) or perform electrical modes of sensing (optoelectrical, electrical, electrochemical, and electromechanical) which are critical capabilities to enable advanced physiological monitoring.

In this work, we demonstrate a minimally invasive implant with a transient geometry that can be inserted subcutaneously and measures important biochemical and biophysical targets such as temperature, cardiac and respiratory activity, pH, and lithium levels. “MiFi" (Minimally invasive Foldable bioelectronic implant) is a multi-modal sensing platform that exploits rapid low-cost fabrication techniques and origami-inspired, foldable planar designs to miniaturize the device up to six times its original length, minimizing the impact of implantation and removal (**Figure 1**). In its unfolded state, the device is compact, thin, and lightweight with planar dimensions of 21 x 21 x 0.3 mm and a weight of 0.24 g. The combination of off-the-shelf components and rapid fabrication simplifies configuration of the MiFi designs, such that sensors can be easily swapped to create a customized monitoring platform for different applications. We validated the MiFi platform extensively in *in vitro*, *ex vivo* and *in vivo* animal experiments to demonstrate the key concepts concerning implantation, operation and removal of the implant.

**Figure 1:**
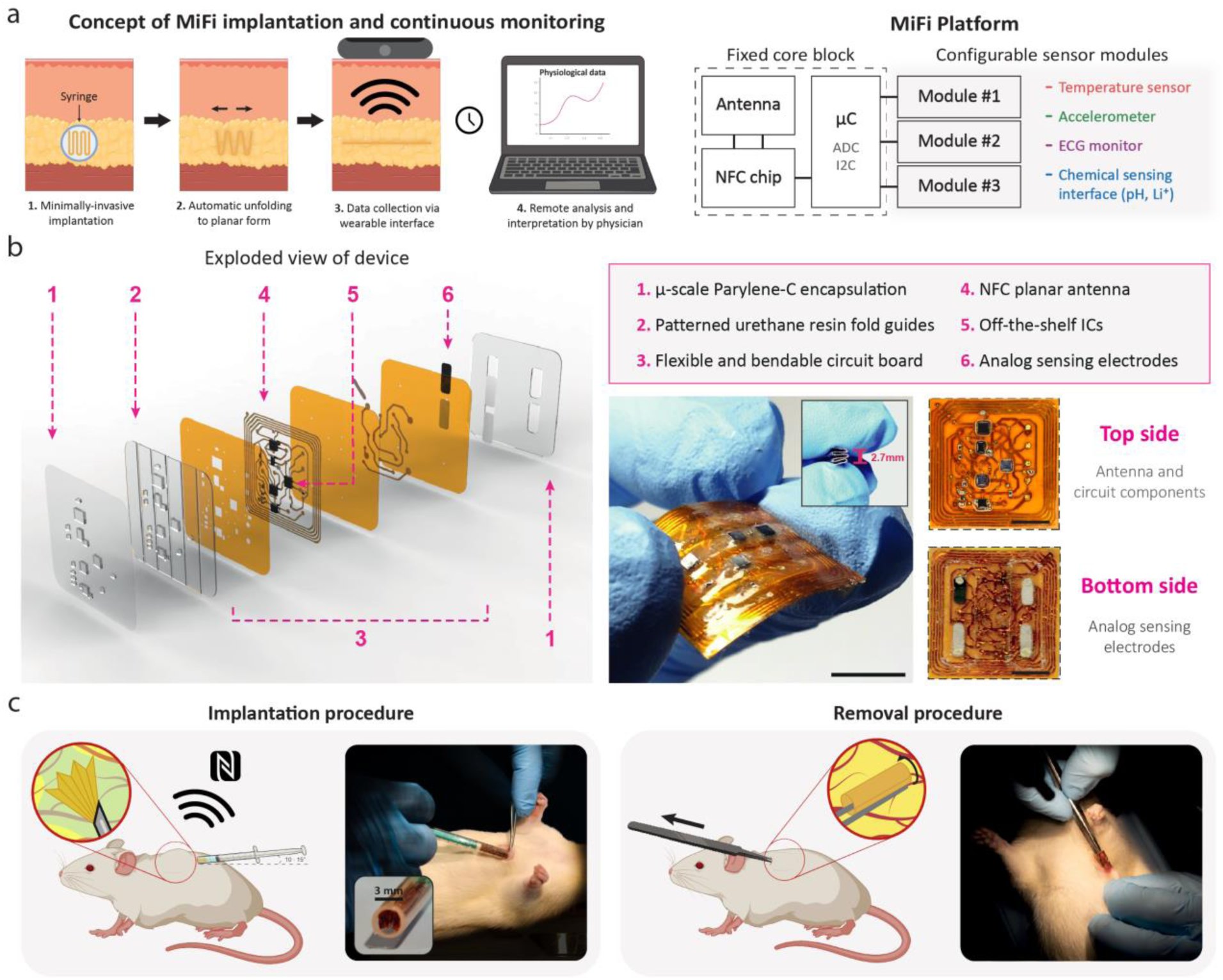
Overview of design and key concepts of a minimally invasive foldable bioelectronic implant (MiFi) for subcutaneous implantation. (a) Conceptual illustration of minimally invasive insertion procedure and continuous operation (left) and schematic of core and sensing functional blocks within MiFi circuit (right). (b) Exploded view rendering of MiFi (left) and photographs of MiFi in folded and planar states (from top and bottom). (c) Conceptual illustration of subcutaneous insertion (left) and removal (right) in rat model with photographs of *in vivo* demonstration.

## Results

### Design

Our design philosophy aims to produce a simple and low-cost sensing platform (MiFi) that is compact during implantation to reduce invasiveness and sufficiently large after implantation to provide the surface area for multiplexed sensing and power/data exchange. The device is designed to be completely foldable along guide grooves to enable precise folding into a compact form for insertion. Once implanted under the skin, in its unfolded form, the MiFi sits flat and seamless underneath the skin, visually undetectable to the naked eye. Figure 1a describes the key features of the MiFi platform. The electronic circuitry is composed of a fixed block that houses the core electronics (and antenna) necessary for power delivery and communication and a configurable sensor block that can be customized to capture up to three different sensing modalities. In this way, the MiFi is designed as a versatile monitoring platform, in which the number and types of sensing modalities can be exchanged to align with a specific application. To reduce development time and costs, the MiFi relies on easily accessible prototyping techniques such as laser cutting to realize modifications in circuit design within hours, speeding up iterations compared to ASIC design and integration.

Fig 1b illustrates an exploded view diagram of the MiFi platform highlighting the three principal design aspects: (i) a custom foldable circuit board manufactured using agile and readily-accessible fabrication methods; (ii) a multi-layered semi-flexible encapsulation designed to facilitate folding of the device and provide electrical passivation; (iii) screen-printed electrodes functionalized for electrical and electrochemical sensing. We embed an NFC (Near Field Communication) planar antenna into the flexible circuit to enable passive powering and wireless communication with a nearby reader (for example, smartphone, computer) without the need for bulky batteries and transcutaneous wires^27–29^. Prioritizing a thin profile facilitates integration with the tissue surface: this not only reduces stress on the surrounding tissue but expands the potential sites for implantation. The larger surface area also prevents migration of the implant, which is a common phenomenon, in highly miniaturized implants^30^. To enable insertion, the circuit must be flexible yet durable enough to withstand the extreme deformations induced by folding. To achieve this, we rely on the inherent elasticity of polyimide (PI)– typically used to secure components during soldering – as the substrate of the circuit, which provides autonomous unfolding of the MiFi back to its planar form after implantation. To facilitate unfolding without additional intervention, we implemented a simple folding design – the accordion fold – as the high tension between the skin and underlying tissue creates little room for multi-plane dynamics. We chose an accordion fold as it provides the most miniaturization in width (the most impactful dimension on invasiveness) while minimizing resistance to unfolding.

To minimize development time and expand the versatility of the device, the MiFi uses only miniaturized off-the-shelf components (max. dimension 1.55 mm). Although smaller than most off-the-shelf integrated circuits (ICs), these components are still large enough to fail mechanically (that is, the component itself or the solder joints) if directly exposed to forces induced by deformation. For this reason, we used a multi-layer encapsulation patterned to facilitate folding while protecting the circuit at extreme deformations (**Fig 1b**). Specifically, we designed columns of semi-flexible urethane resin to stabilize components and their connections, while the guide grooves between columns remain fully bendable. This concept facilitates folding in two primary ways: (i) by guiding the fold to occur at a precise location (that is, not underneath any of the components); (ii) by creating weak points to concentrate folding-induced stresses and further protect the surrounding components. Finally, a micron-scale (7 µm) film of parylene-C electrically passivates the circuit from the surrounding tissue while still preserving the flexibility of the substrate.

To demonstrate the versatility of the MiFi, we developed three configurations of the MiFi platform that span four different sensing modalities: temperature, electrocardiography (ECG), electrochemical sensing, and inertial sensing. We chose a temperature sensor, ECG monitor, and inertial sensor (heart rate, breathing rate) for their overwhelming utility in diagnostic monitoring as three of the five vital sign measurements^31^. We also demonstrate functionality for two potentiometric electrochemical sensors – pH and lithium (Li^+^) – as a proof-of-concept for more advanced chemical monitoring. While pH is used as a clinical indicator (for example, infection or inflammation), the same electrodes can also be functionalized with the appropriate molecules to detect other clinically relevant markers, including metabolites, antibiotics, and alcohols^32–35^.

All sensor electrodes were screen-printed onto a PI substrate and attached to the MiFi as a modular unit. Because many electrochemical sensors utilize similar sensing circuitry (in this case, open circuit potentiometry), we can easily modify electrodes for other electrochemical sensing applications without the need for redesign and refabrication. In this regard, we chose to integrate a Li^+^ ion sensor because of its prominence as the primary therapeutic treatment for bipolar disorder (BPD): a life-threatening chronic mental illness with a narrow and highly sensitive therapeutic range (∼0.5 mmolL^-1^) which is toxic at larger concentrations^36^.

### Fabrication and characterization of a foldable electronic platform

#### Flexible circuit board

NFC protocol requires 13.56 MHz for data exchange: hence, it is important to investigate how folding of a flexible circuit board for miniaturization would affect the performance of a planar NFC antenna. We first measured the reflection coefficient (S11) of a set of commercially produced flexible antennas (model: flexPCB, manufacturer: PCBWay, Shenzhen, China) between 11 – 17 MHz (**Fig 2a**). S11 is a standard parameter that provides information concerning the optimum operating frequency (resonant frequency) of the antenna.

**Figure 2:**
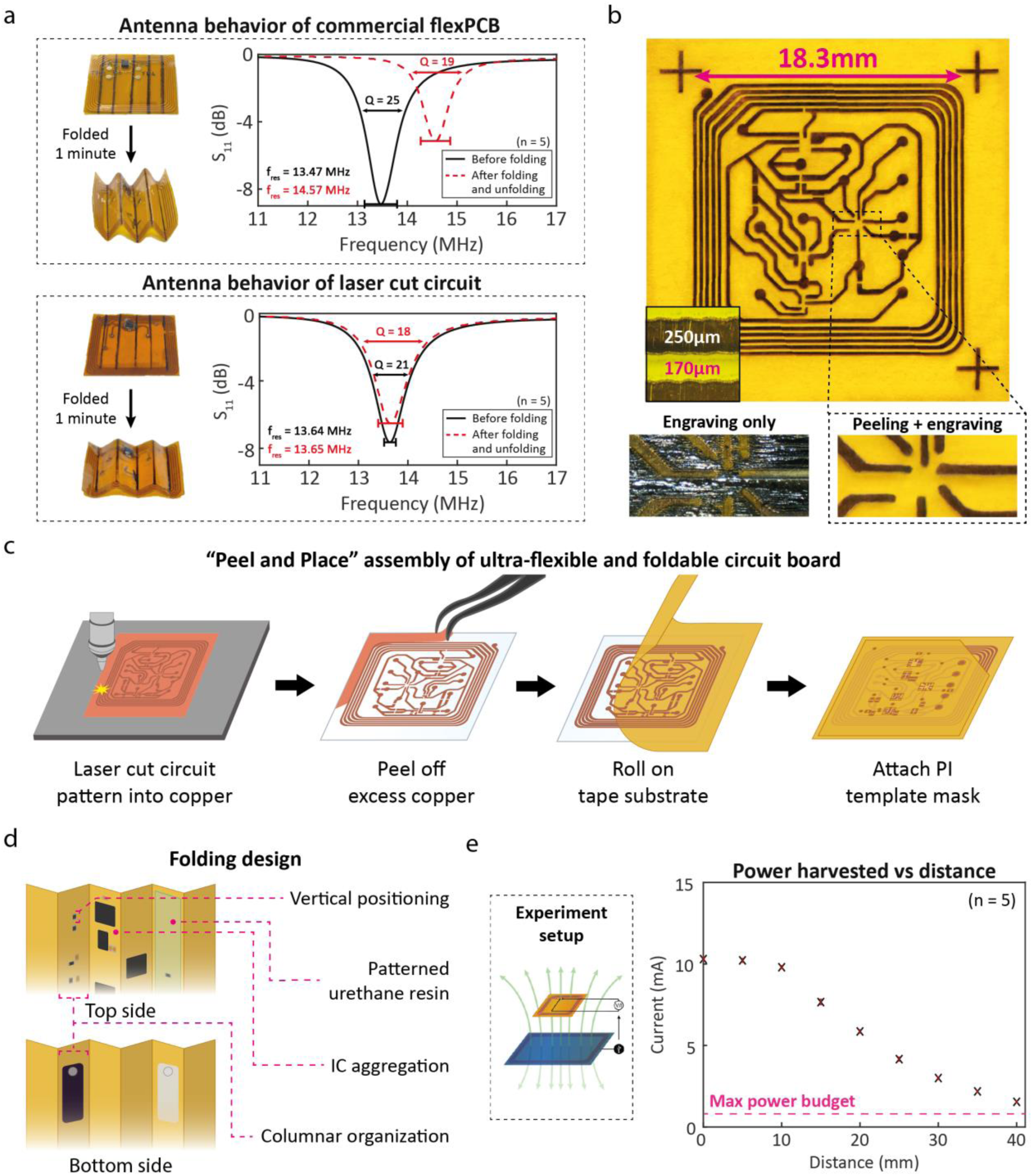
Fabrication, design, and characterization of foldable antenna and circuit board. (a) Comparison of NFC antenna tuning (S11) using in-house Peel and Place Assembly (PPA) fabrication of circuit board (185 µm) and commercially manufactured flexible circuit board (80 µm) after folding for 1 minute and resting for 5 minutes. (b) Photograph of single layer of PPA- fabricated circuit (top), trace patterning resolution by only engraving copper foil (left) versus peeling laser-cut sections of copper and engraving densely connected areas (right). (c) Illustration of PPA method for a single circuit layer. (d) Illustration of design layout strategies used to enable folding of a populated PPA-fabricated circuit board without causing damage to components. (e) Change in magnitude of power harvested from NFC (mA) with increasing distance between planar NFC antenna and commercial reader (illustrated), compared against the maximum power budget of all three MiFi circuit configurations (780 µA).

To maximize flexibility of the circuit board, we chose the thinnest substrate available for purchase (80 µm). In this experiment, we folded each antenna in an accordion pattern and maintained the fold for one minute before allowing them to unfold. After five minutes, we remeasured S11 over the same range of frequencies. We observed a marked shift in resonant frequency of approximately 1.1 MHz. We also noted that the flexPCB was unable to recover its unfolded geometry due to the stiffness of the materials used by the manufacturer. Commercially available flexPCBs would, therefore, not be suitable for applications requiring transient geometries that require folding such as minimally invasive implants.

To satisfy the requirements of an ultra-flexible and foldable (bending radius < 1 mm) circuit board, we developed our own circuit board fabrication method (“Peel and Place Assembly” – PPA) using specialized materials and easily accessible tools that can be found in most academic laboratories. We used a 70 µm-thick PI tape (5413) as the substrate for the circuit due to its high elasticity relative to commercial-grade PI flexPCBs. We used a standard fiber laser cutter (20W, Trotec Speedy 300) to define copper traces at high precision (∼50 µm), which enables the integration of even the smallest ICs (for example, WLCSP) and components (**Fig 2b**). To improve the resolution around densely packed and narrow traces, we initially assessed engraving as an alternative method of patterning the copper film. Patterning this way, however, left thin fibers of copper along edges (compromising electrical isolation), increased fabrication time (from approx. 2 to 40 minutes per layer), and prevented transfer to the PI substrate: this is particularly detrimental to the antenna, where even slight deviations in positioning impact its electromagnetic properties. For these reasons, we patterned most traces by laser cutting a thin copper foil (10 µm) while particularly intricate areas (for example, pinout of WLCSP ICs) were engraved at a higher intensity for enhanced resolution (see SI: Fig S2 for patterning layout). We used adhesive films during circuit assembly to easily transfer and place copper traces on the PI substrate, peeling away the excess copper to reveal the final circuit pattern (**Fig 2c**). Unlike traditional subtractive methods of patterning such as chemical etching, our method eliminates the need for clean-room facilities, hazardous waste management, and time-consuming cycles of fabrication. To fabricate multi-layer circuit boards, the adhesive substrate layers are aligned and stuck together. Electrical continuity was achieved through soldered via points (see Methods for further information). Thus, we created multi-layer connections using simple and well-established commercial practices without the need for expensive and laborious microfabrication techniques. To improve the mechanical stability of on-board components, minimize the stresses on components during deformation, and to aid with precise placement of components during population, we created a component template mask by laser cutting a thin sheet of PI (25 µm thickness, Kapton HN) which is adhered on the top layer of the circuit (SI: Fig S3).

We repeated the folding experiment using the same antenna design patterned on PPA- fabricated circuit boards to validate the effects of increased flexibility on antenna performance (**Fig 2d**). The resonant frequency of the planar NFC antenna remained practically unchanged (approx. 10 kHz shift) while the circuit board achieved a visibly higher degree of recovery towards its planar form. This is because highly flexible substrate and trace materials minimize the occurrence of sharp creasing during folding: creasing deteriorates the electromagnetic (via detuning) and mechanical (via permanent deformation) behaviors of the MiFi. This qualitative observation was confirmed in a series of experiments performed using a programmable tensile and compression tester (MultiTest 2.5-i – Mecmesin). The commercial flexPCB exhibited greater mechanical hysteresis upon folding and required a greater peak compressive force (2.3 N vs 1.3 N) to achieve a bending radius of 1 mm despite being less than half as thick (80 vs 185 microns) in comparison to the PPA-fabricated circuit board (see SI: Fig S4 for more details on mechanical characterization).

#### Circuit layout

To minimize the potential for damaging interconnects, solder joints and components during folding, we arranged components into six columns to align with the accordion folding pattern (**Fig 2d**). All components and vias were positioned in the center to minimize impact on the solder joints. We also stabilized particularly miniaturized passive components (max dimension: 0.6mm) by favoring vertical alignment when possible and positioning them next to larger and more stable ICs. Finally, we optimized positioning of ICs to prevent stacking or collisions that jeopardize mechanical stability and minimize the thickness of the overall device when folded (see SI for more detail).

#### Encapsulation

To guide folding into an accordion pattern we created guide groves by depositing a semi-flexible urethane resin (TASK™ 3 – Smooth-On UK) in the form of columns. We chose TASK™ 3 because of its excellent adhesive properties and flexibility (tensile modulus ∼100x higher than epoxies traditionally used to stabilize circuit components). Maintaining flexibility is important for improving the contact between the electrodes and tissue while minimizing the foreign body response to the implant^37^. To improve biocompatibility and waterproofing, we also deposited a conformal film (thickness: 7 µm) of Parylene-C via chemical vapor deposition. Films thinner than 7 µm were less stable under folding and compromized waterproofing while thicker layers were too rigid to induce unfolding following injection.

#### Power characterization of MiFi

Because NFC requires proximity between a receiver and transmitter, we also characterized the efficiency of power delivery to the MiFi antenna as a function of vertical distance (**Fig 2e**). Since the MiFi is designed to function inside the body, it is naturally separated from the reader: hence, delivery of power over a range of distances is an important parameter. In this experiment, we used the configuration of the MiFi with the largest power requirement (ECG, pH and temperature sensing), requiring 780 µA. The power harvested remained well over the maximum power required (780 µA) up to 30 mm of separation. Beyond 30 mm, sufficient power is still available to maintain electrical operation, however, communication is lost due to a loss of efficiency during amplitude modulation (needed to encode data for transmission). Although communication efficiency can be improved further with the use of an impedance matching circuit (which would require additional footprint on the substrate), for subcutaneous implantation, 30 mm is sufficiently large for reliable operation.

### Design, fabrication, and characterization of configurable sensor modules

#### Chemical sensing interface

We designed a chemical sensing interface (CSI) to enable simple, low-power potentiometric sensing of pH and Li^+^ within the interstitial fluid (ISF). The simplicity of the sensing principle keeps the footprint of the circuitry small: we incorporated a buffer to prevent current flow from the core computing block (current flow can shift the potential) and a reference voltage (Vref = Vcc/2) to capture negative shifts in equilibrium potential (**Fig 3a**). To sense pH, we electrodeposited a thin film of polyaniline (PANI), a pH sensitive polymer, on a flexible carbon electrode. We chose to use PANI because of its easy processability, excellent biocompatibility, and linear sensitivity^38^. For Li^+^ monitoring, we adapted the protocol developed by Amirghasemi et al. to create a Li^+^-sensitive membrane (containing Li^+^ ionophore VI) that we screen-printed onto a flexible carbon electrode^39^. Finally, we screen-printed Ag/AgCl ink to create a flexible electrode that acts as a pseudo-reference. We conducted a set of calibration experiments for each electrode to characterize their sensitivity, linearity, and stability in phosphate buffer solution (PBS) as it mimics the composition of ISF (**Fig 3b)**. The electrodes produced a high degree of linearity (pH: R^2^ = 0.98, Li^+^: R^2^ = 0.99) and sufficient sensitivity for digitization (pH = -74 mV/pH, Li^+^: 43.1 mV/dec) that were not affected by integration into the MiFi platform and exhibited low drift (pH: 0.21mV/hr, Li^+^: 0.28 mV/hr) when monitoring over longer durations (see SI for further characterization of each sensor). The Li^+^ electrode achieves higher sensitivity (55.1 mV/dec) and micromolar limit of detection (LOD, < 10 µM) in non-ionic solutions such as TRIS (SI: Fig S7). Cross-interference from primarily sodium within PBS (and ISF), however, reduces selectivity to Li, particularly impacting the LOD (0.32 mM)^39^.

**Figure 3:**
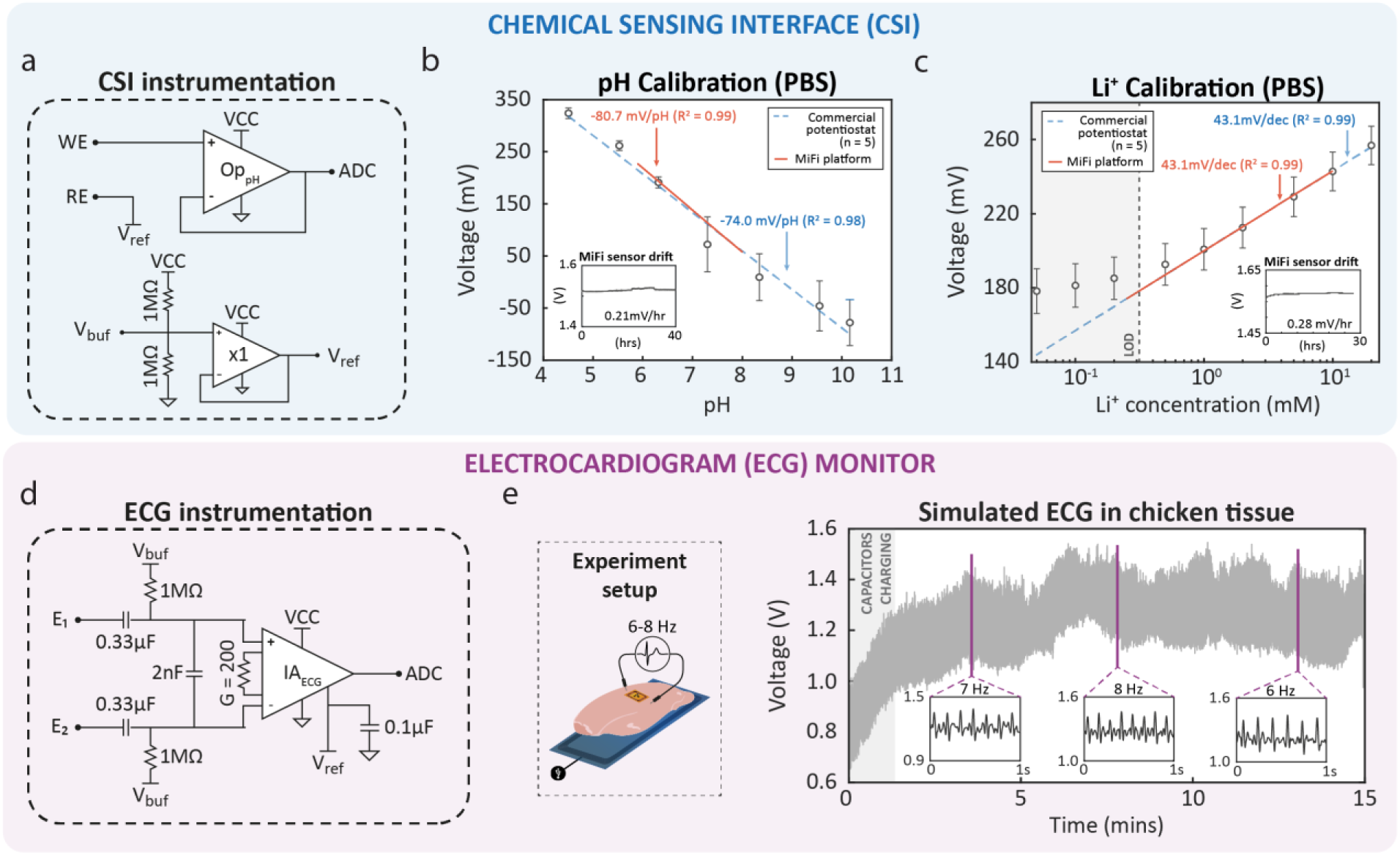
**Design and characterization of ECG monitor and chemical sensing interface**. (a) Schematic of analog front end for potentiometric chemical sensing interface. (b) Calibration curve of PANI-based pH sensor when connected to commercial potentiostat (using standard double-junction Ag/AgCl reference electrode, n = 5) and MiFi circuit (using screen printed Ag/AgCl ink reference electrode). Inset: stability of pH sensor connected to MiFi circuit. (c) Calibration curve of Li^+^ ionophore-based Li^+^ sensor when connected to commercial potentiostat (using standard double-junction Ag/AgCl reference electrode, n = 5) and MiFi circuit (using screen printed Ag/AgCl ink reference electrode). Inset: stability of Li^+^ sensor connected to MiFi circuit. (d) Schematic of analog front end for two-electrode ECG monitor. (e) *Ex vivo* experiment of simulated ECG monitoring using signal-generated ECG waveform (6 - 8 Hz) transmitted across chicken tissue (setup illustrated).

#### ECG monitor

We used a miniaturized instrumentation amplifier (IA; MAX414000) to enable two-electrode biopotential monitoring across the heart (**Fig 3c**). To minimize the footprint of the ECG front-end, we integrated a high-pass filter (fc = 0.007 Hz, pulled up to Vbuf) prior to amplification to minimize the effect of saturation from localized potential drifts (see SI: Fig S8 for filter design optimization). Given the limited memory of the µC and communication protocol set by our NFC chip (NTAG I2C plus), we were able to achieve a sampling rate of 334.5 Hz (the fastest wave of ECG in rats is roughly 10 ms in duration). For the sensing of cardiac biopotentials, we used screen-printed Ag/AgCl electrodes because of their high conductivity and low polarizability in ionic solutions, both of which are crucial for ECG monitoring^40^. To ensure adequate capture of the difference in potential during cardiac depolarization, we distanced the ECG electrodes as far from each other as possible (∼10 mm). We characterized the performance of the ECG circuit design using chicken tissue as a simulated biological monitoring environment (**Fig 3d**). The amplified output of the ECG monitor exceeded the resolution of the ADC (∼3mV) by over 80x and was able to clearly capture a range of physiologically relevant heart rates (6-8 Hz in rats). Due to the capacitance of the high pass filter, we observed a short transient charging period (∼90 seconds) before ECG output stabilized (see SI for further details about circuit design).

### Digital sen3sing modules

We used a digital silicon bandgap temperature sensor (AS6212) that monitors subcutaneous temperature with high-accuracy (± 0.2 °C) and calibration-free operation at a miniaturized scale (< 1.5 mm). Unlike commercial thermocouples, the milli-degree precision resolution of the AS6212 makes the MiFi well-suited to biological applications where temperature changes are more tightly regulated (< 1 ° C). To measure heart rate and breathing rate simultaneously from the chest, we used a digital three-axis MEMS accelerometer (BMA530)^41,42^. Unlike subcutaneous ECG, inertial heart rate monitoring does not require any electrodes, so does not face issues related to biofouling, improper electrode-tissue contact, and electrode degradation over time. By using only one axis to monitor cardiac and respiratory movement, we achieved a sampling rate of 235 Hz for inertial monitoring.

#### Multiplexed sensing

The MiFi platform is highly reconfigurable and can be used for multiplexed *in vivo* sensing with up to four sensors. To demonstrate this concept, we produced versions of MiFi with three different configurations for multiplexed sensing. The first configuration (Config 1) measured subcutaneous temperature, pH, and ECG monitoring **(Fig 4a**). The second configuration (Config 2) measured subcutaneous temperature, interstitial pH, heartrate, and breathing rate monitoring **(Fig 4b**). By replacing the ECG monitor with the BMA530, we expand the diagnostic capability of the device while simultaneously reducing circuit complexity. The third configuration (Config 3) acts as an electrochemical unit and measures interstitial pH and Li^+^ concentration **(Fig 4c**).

**Figure 4:**
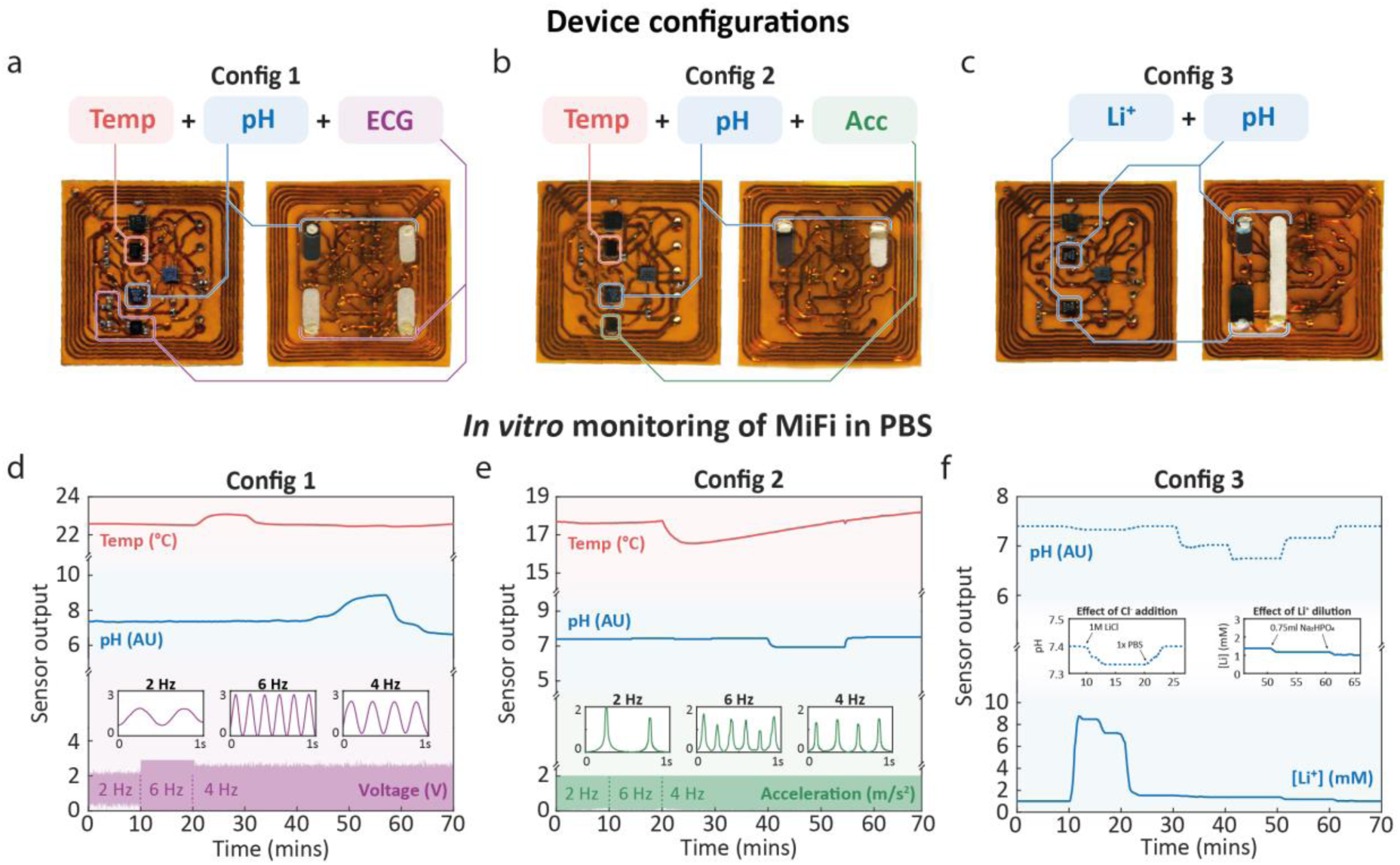
Configuration and *in vitro* validation of multiplexed monitoring. (a) Photographs of Config 1 of MiFi monitoring temperature, pH, and ECG. (b) Photographs of Config 2 of MiFi monitoring temperature, pH, and acceleration (for heart and breathing rate detection). (c) Photographs of Config 3 of MiFi monitoring pH and Li^+^. (d) *In vitro* experiment in PBS to validate multiplexed monitoring of Config 1 in response to changes in temperature, pH, and simulated biopotential in PBS. (e) *In vitro* experiment in PBS to validate multiplexed monitoring of Config 2 in response to changes in temperature, pH, and simulated motion. (f) *In vitro* experiment in PBS to validate multiplexed monitoring of Config 3 in response to changes in pH and Li^+^ concentration.

To demonstrate successful monitoring as a multiplexed platform, we characterized each configuration *in vitro* in PBS. We generated a stimulus for each sensor module (for example, increase in temperature, decrease in pH) to validate successful operation of the MiFi, confirm the absence of cross-interference between sensors (this is especially important when more than one analog sensor is integrated), and to verify waterproofing of the Parylene-C encapsulation (specifically after folding). We observed successful monitoring of changes in temperature, pH, ECG, and acceleration without generating cross-interference or instability in the other sensors (**Fig 4d,e**). Config 3 exhibited cross-reactivity between the pH and Li^+^ sensors, however, these artefacts were attributed to indirect changes in solution composition: (i) spiking the solution with LiCl generated a slight bias in the Ag/AgCl pseudo-reference electrode which shifted pH; (ii) modifying the pH without changing solutions increased the volume and diluted the concentration of Li^+^ (see Methods for further details).

### Real-time *in-vivo* monitoring in anesthetized rats

To test the functionality of the MiFi in living animals, we performed a series of terminal experiments using anesthetized rats (Sprague Dawley: male, 368-719 g). The procedure included implantation, monitoring and removal (postmortem) of the MiFi. Where possible, we also used standardized laboratory equipment for comparison (such as ECG) to benchmark the functions available within MiFi.

### Implantation procedure

The MiFi was folded and placed inside of a 1 ml syringe (tip cut off) which acted as the injector. To prepare the area for implantation, a small section of fur was shaved and sanitized. A small incision of approximately 3 mm was created to allow for entry of the injector (**Fig 5a**). Due to the layer of fascia (connective tissue) present in rats that sits between the skin and muscle (not present in humans), a closed pair of scissors was used to separate the fascia layer while minimizing trauma to the surrounding tissue. Once a pocket was formed, the syringe inserted the folded MiFi into the subcutaneous space. As the device left the syringe, it immediately unfolded such that once injection was complete, the MiFi had already reached its planar form and was not visible as it sat underneath the skin (see SI for further details).

**Figure 5:**
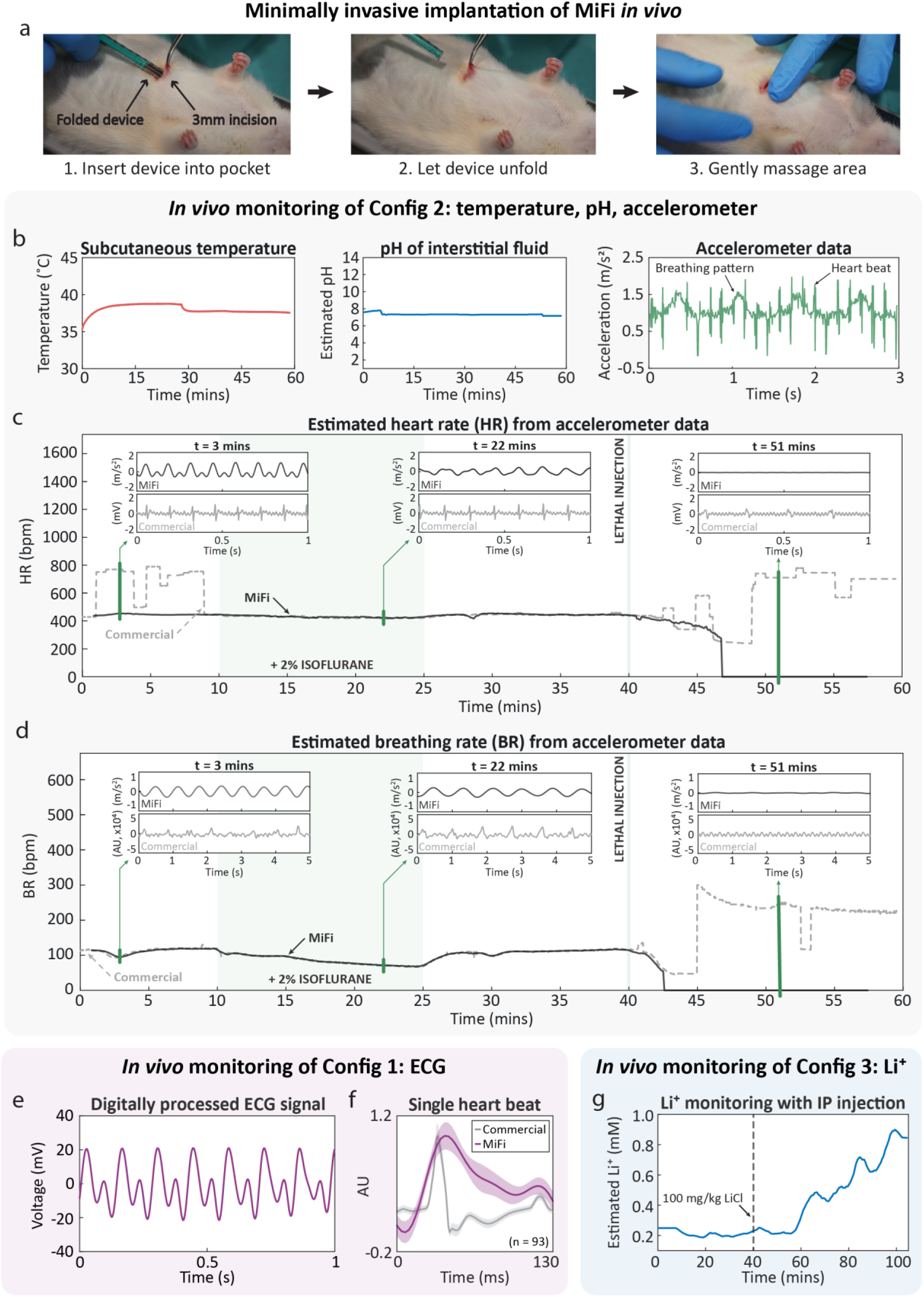
***In vivo* minimally invasive implantation and continuous operation in live rats.** (a) Photographs of minimally invasive insertion procedure of folded MiFi into subcutaneous space of anaesthetized rats. (b) Recorded subcutaneous temperature (left), pH of interstitial fluid (middle), and accelerometer data (right) from *in vivo* implantation and operation of Config 2. (c) Estimated heart rate and breathing rate (d) from accelerometer data over multiple interventions (changes in level of anesthetic, lethal injection) in comparison to heart and breathing rate data collected from commercial small animal monitoring system. Insets compare the filtered cardiac and respiratory signals between the MiFi and commercial monitoring system at various time points. (e) Recorded ECG signal (digitally processed using FFT) from *in vivo* implantation and operation of Config 1. (f) Comparison of the average ECG heartbeat recorded from MiFi and commercial monitoring system (n = 93). (g) Recorded Li^+^ concentration of interstitial fluid from *in vivo* implantation and operation of Config 3. At t = 40 minutes, 100mg/kg LiCl is administered to animal subject through intra-peritoneal injection.

### *In vivo* monitoring of subcutaneous temperature, interstitial pH, heart rate, and breathing rate (Config 2)

A live urethane-anaesthetized rat was monitored in a resting state for approximately ten minutes, after which 2% isoflurane was administered by mask delivery. We expected that deepening the anaesthetized state of the animal would reduce the respiration rate and indirectly reduce the heart rate. This physiological state was maintained for a further fifteen minutes, after which the isoflurane supply was removed to recover breathing and heart rate. The animal was left in this state for fifteen minutes. The animal was then culled using an intraperitoneal (IP) injection of pentobarbital (2 ml). We maintained monitoring of heart rate and breathing rate during this period to capture the decline and consequent cessation of heart and breathing patterns.

After folding and implantation, all three sensors remained functional with no indication of component detachment or water penetration. The temperature sensor continuously monitored the subcutaneous temperature, regulated to ∼38 °C by a heating pad positioned underneath the animal throughout the experiment (**Fig 5b** – left). After a short conditioning period (under 10 minutes, from dry to wet), the interstitial pH stabilized to pH 7.3 throughout the duration of live monitoring and declined to pH 7.0 after 53 minutes (**Fig 5b** – middle). We believe that the increase in acidity, recorded approximately 13 minutes after lethal injection, is indicative of the buildup of acidic waste (for example, carbonic acid) that occurs as blood circulation stops, as also reported by others^43,44^.

The accelerometer output captured the breathing and heart signals over the duration of the experiment: high frequency spikes representing the heart pattern were modulated by lower frequency oscillations which correspond to the breathing pattern (**Fig 5b** – right). We applied a series of digital filtering techniques (described in Methods) to isolate the breathing and heart patterns. From these patterns, we also calculated the heart and breathing rates and compared them against measurements taken by a non-invasive commercial monitoring system (Harvard Apparatus, **Fig 5c**).

Over one hour of recording, the MiFi continuously captured the breathing and heart rate of the animal while the commercial system demonstrated long periods – up to 10 minutes prior to culling – of malfunctioning heart rate estimation (unstable and far outside of physiological range). The commercial accuracy degraded even further post-culling, when both heart and breathing rate estimation remain elevated. Because the MiFi uses motion as its sensing modality, it is completely immune to biologically induced electrical noise (which leads to erroneous results when biopotentials are solely used to measure cardiac activity). The subcutaneous placement of the MiFi also insulates the sensors from non-physiological motion artefacts that typically affect non-invasive devices (for example, sensor sliding against fur/skin), improving their stability during continuous monitoring. Excluding the regions of malfunctioning, the MiFi exhibited a mean difference of less than one bpm (-0.85 bpm ± 9.6) for breathing rate estimation and 2.57 bpm (± 5.25) for heart rate prediction.

Analyzing the heart pattern under homeostatic conditions, we observed that each heart pattern was characterized by two peaks: a large prominent peak – corresponding to aortic opening – followed shortly after by a smaller peak – corresponding to aortic closing (**Fig 5c** – left inset)^45^. After administering isoflurane, we recorded a decline in heart rate of ∼24 bpm as expected. We also noticed additional modulation in the signal pattern: the ventricular peak decreased in prominence by 43% and the atrial peak was no longer detectable, providing further insight into the physiology of the animal in real-time as chamber contraction weakened from elevated anesthetization (**Fig 5c** – middle inset). Because ECG only captures electrical impulses, the ECG signal remains unchanged and is unable to encode this additional physiological information. Non-temporal modulation of ECG was only observed under extreme changes, over ten minutes after lethal injection was administered (**Fig 5c** – right inset). The breathing pattern captured a similar but more severe decline in breathing rate (119 bpm to 66 bpm) after administering isoflurane (**Fig 5d**). Despite the MiFi and commercial monitor both using motion to detect respiration, the MiFi exhibited a higher prominence in amplitude modulation: the RMS (root mean square: used to describe the average power of a signal) of the commercial signal is only ∼59% of the RMS of the MiFi signal (when normalized for range). We noted that the MiFi was additionally less prone to high-frequency artefacts, particularly noticeable after culling (**Fig 5d** – right inset). The visibly higher signal-to-nose ratio demonstrated by the MiFi through inertial sensing ensures robust peak detection during breathing rate estimation and consequently highlights the value of monitoring underneath the skin.

### In vivo monitoring of ECG

We conducted preliminary *in vivo* experiments and discovered the cutoff frequency (fc = 0.007 Hz) was too low to suppress low-frequency drifts from surrounding electrical activity (for example, by skeletal muscle). Thus, to maximize the quality of the input biopotential signal during *in vivo* operation, we modified the circuitry to increase the cutoff frequency (fc = 0.48 Hz). We validated the ECG monitor *in vivo* as part of Config 1, which also integrates temperature and interstitial pH monitoring (See SI for the results on temperature and pH for this experiment). Despite successful monitoring in simulated conditions *ex vivo*, we were unable to replicate the same signal quality in the raw ECG data. The ECG signal was mostly composed of high frequency noise, likely from surrounding skeletal muscle. Inspecting the frequency spectrum of the data, however, we observed a strong peak at 7 Hz (equal to 420 bpm), indicating that the ECG waveform was still captured (SI: Fig S14). Thus, we employed Fast Fourier Transform (FFT) filtering to isolate the physiologically relevant frequency components and reconstruct the ECG waveform (**Fig 5e**). From the reconstructed signal, we identified two characteristic peaks: a high amplitude peak resembling the QRS waveform followed by a low amplitude peak resembling the P waveform. We overlaid our ECG signal with the average ECG waveform recorded by the commercial monitor to validate our findings (**Fig 5f**). The two signals clearly aligned, though the reconstruct ECG waveform was unable to provide the same quality of information (for example, T wave lost) due to aggressive filtering in the frequency domain.

### In vivo monitoring of lithium within the interstitial fluid

We conducted a third *in vivo* experiment using Config 3 to validate the performance of the electrochemical Li^+^ sensor. Since Li^+^ is not a localized biomarker, we inserted the MiFi at the scruff of the animal to facilitate implantation (SI: Fig S10). We first monitored the live anesthetized rat in a resting state for 40 minutes to establish a reference baseline (after an initial 30-minute waiting period to allow the sensor to equilibrate after implantation). We then administered a dose of LiCl (100mg/kg of bodyweight) through IP injection to allow the Li^+^ to diffuse into the bloodstream and reach the sensor naturally via ISF partitioning (**Fig 5g**)^46^. This method of injection verified that the sensor was able to detect true interstitial levels of Li^+^, as opposed to direct injection of LiCl subcutaneously underneath the site of implantation. Post-injection, we continued monitoring for one hour to allow sufficient time for the Li^+^ dose to fully partition into the ISF (∼45 minutes) and to comply with the animal license (long-term and recovery experiments were not permitted)^47^.

To improve the stability of the output of the Li^+^ sensor and compensate for fluctuations in the potential of the pseudo-reference electrode (chloride from LiCl affects stability of Ag/AgCl), we performed differential analysis by subtracting artifacts identified in the potentiometric output of the pH sensor (undergoing no intervention in this experiment) from the Li^+^ sensor data (see SI: Fig S11). We then converted the cleaned potentiometric signal to estimated Li^+^ concentration using the calibration curve presented in Fig 3c (see Methods for details). We observed a baseline of approx. 0.25 mM of Li^+^ prior to injection of LiCl. Although animals accumulate small amounts of Li+ in the body through the consumption of various food and water, the elevated baseline is likely due to the cross-interference of sodium ions present in the ISF^48^. Post-injection, the signal remained unchanged for approximately 20 minutes (in line with reported extracellular partitioning times of Li^+^) before the concentration of Li^+^ increased above the LOD of the sensor (0.32 mM)^47^. The Li^+^ concentration continued to increase over the duration of the experiment, reaching a concentration of 0.9 mM in the ISF one hour after administration.

## Discussion

The MiFi platform, as presented here, enables minimally invasive subcutaneous implantation of bioelectronic sensors for biophysical and chemical monitoring. Despite its relatively large surface area and dense component population, the MiFi can be implanted using minimally invasive procedures similar to ICMs, which are suture-free and take less than 15 minutes to implant^49^. Our custom PPA fabrication method creates a new class of robust foldable electronics that solves a critical design challenge in the development of implants with transient geometries to minimize invasiveness. Our use of agile fabrication technologies such as laser cutting and substrate-based patterning remains compatible with roll-2-roll processes that facilitate scaling up to mass manufacturing. By avoiding chemical etching for circuit patterning, solid copper waste accumulated from peeling can also be easily recycled, improving sustainability and environmental impact of electronics. Depending on MiFi configuration, one device costs between US $4-7 to manufacture (when ordering in bulk, see SI for price breakdown): the largest cost comes from the integration of multiple IC components, which can be further reduced by transitioning to ASIC-based sensor design (though time and cost of development will increase).

We demonstrated successful implantation and operation of the three developed configurations of the MiFi (including temperature, CSI, ECG, accelerometer) through *in vivo* experiments in anaesthetized rats as proof-of-concept. We believe the degradation in the performance of the ECG sensor was attributed to oversimplification of the front end, which we prioritized to ensure that the sensing circuitry fit within the limited available area. Including a low-pass filter to attenuate high-frequency artefacts and adding an additional amplification stage (for example, 5x) would likely improve the quality of the signal in future experiments. Though biophysical measures (for example, heart and breathing rate) could be validated with commercial instruments, we did not have the instruments available to monitor pH and Li^+^ levels continuously in the ISF: this limitation can be addressed using other established laboratory techniques in more extensive animal studies. While the physiological range of pH in the ISF is relatively well established, the interactions of Li^+^ within various mechanisms in the body are still not completely understood (for example, tissue absorption and active transport mechanisms)^50^. For these reasons, it was not possible to validate the correlation between the peak Li^+^ concentration detected by the MiFi and the IP-administered Li^+^ concentration. We hope that in the future MiFi can help contribute to expanding the understanding of Li^+^ dynamics and equilibration mechanisms throughout the body.

The MiFi technology presented here has three main limitations: (i) Despite the inherent flexibility and low cost of ink-based electrodes, they lacked the stability and mechanical robustness needed for long-term high-quality continuous monitoring *in vivo*. Thin film deposition techniques (for example, physical vapor/atomic layer deposition) can instead be used to produce metal-based electrodes (for example, titanium, palladium, gold) that offer superior electrical and mechanical stability at a relatively low cost^51,52^. (ii) While circuit and electrode patterning were executed using automated fabrication tools, we carried out the majority of assembly manually, which compromized the repeatability of fabrication. Given that these manual processes are still in line with high-volume, commercial manufacturing processes, however, they can be outsourced to manufacturing companies for automated production. (iii) Using off-the-shelf components increased the minimum width (and height) of each folding plane. This can be addressed by using bare dies rather than packaged components, which can further reduce the folded dimensions of MiFi and allow for even greater minimization during implantation.

Due to existing limitations with the longevity and stability of biosensors, the MiFi is best suited as a short-term implant (up to 30 days)^53^. As demonstrated with the Eversense CGM, for example, intermittent replacement implantation procedures can provide greater comfort and improved accuracy compared to current transdermal biosensors^11^. The MiFi can additionally be paired with implantable drug delivery systems to achieve completely automated and optimized treatment, known as ‘theragnostics’ (THERApeutic diaGNOSTICS)^54–56^. As healthcare moves away from generalized health models driven by statistics, the integration of multiplexed physiological monitoring into a single implant, as achieved with the MiFi, can be used to develop a comprehensive personalized health profile of the patient through the implementation of sensor fusion or train AI models, providing unique health insights and warning signs that cannot be extracted using just one biomarker^57–59^.

## Methods

All chemicals, unless stated otherwise, were purchased from Merck. All wireless data transmission was captured using the open-source serial application ‘SerialPlot’ and all data processing was done in MATLAB.

### Device fabrication

#### Double-layer circuit board

We designed the top and bottom layers of the circuit board using Adobe Illustrator and laser-cut (Model: Trotec Speedy 100; Wavelength: 1 µm; Power: 20 W) the designs into two thin sheets of thin Copper (Cu) foil (10µm-thick). We rolled vinyl transfer tape onto the patterned Cu foils as a temporary substrate and peeled away any unused sections of Cu using a pair of tweezers. We then rolled a thin heat-resistant PI tape (70µm-thick, 5413 – 3M) onto the Cu-vinyl substrates and peeled the transfer tape off. We laser-cut the soldering template into a thin PI sheet (25 µm-thick, Kapton HN – 3M) and attached it to the top layer using alignment markers positioned in each corner of the circuit board design. We defined via holes (0.4 mm diameter) by laser-cutting the bottom side of the PI-Cu-PI top layer such that only the PI substrate was ablated, exposing the copper traces at designated locations. We then deposited and melted solder paste (SMD291SNL10 (Sn96.5/Ag3.0/Cu0.5, 217°C) – CHIP QUIK) onto the exposed copper to enable multi-layer connection. We attached the top and bottom layers using the adhesive from the PI substrate of the bottom layer. We heated the combined layers on a hotplate to re-melt the solder bumps and electrically connect the top and bottom layers. We then realized the final dimensions of the circuit board using a guided blade.

#### Component attachment

We soldered all WLCSP ICs to the circuit board using conventional solder reflow on a hotplate with the addition of flux (Soldering flux paste 8341 – MG Chemicals) to facilitate solder melting and wetting onto the copper connections. We soldered the NFC chip and passive components to the circuit board using a low-temperature solder paste (SMDLTLFP10T5 (Sn42/Bi57.6/Ag0.4, 138°C) – CHIP QUIK) and conventional solder reflow on a hotplate. Once all connections were made, we deposited a thin border of flexible epoxy (ET515 - Permabond) around each component and left it to cure at 60°C overnight. See SI for details of device programming and sensor communication.

#### Encapsulation

We cut strips of PI tape (0.4 mm width) using a plotter cutter (CE6000-40 - Graphtec) and adhered them onto the circuit board at the designated guide grooves. We treated the circuit board with oxygen plasma (High Power Expanded Plasma Cleaner (PDC-002-CE) – Harrick Plasma) for 20 minutes, after which we prepared and deposited a urethane resin (0.1mL part A : 0.1mL part B. TASK 3 – Smooth On) between the PI strip masks. Twenty-three minutes after mixing (before pot life runs out), we peeled away the masks and the patterned resin was left to cure at room temperature. The resin was then post-cured at 60°C for at least one hour. Prior to parylene-C encapsulation, we masked the via holes at the bottom of the circuit board (for electrode attachment) using PI tape. We folded the device for approximately five minutes, after which we released the fold and placed the device inside the chamber of a parylene coater (Parylene-C deposition system (2010) – Specialty Coating Systems). Folding the device prior to coating prevented the parylene film from tearing because of folding. To achieve a monolithic coating around the circuit board of appx. seven microns, we used 13.9g of parylene dimer.

Finally, we removed the mask and overlapping parylene-C film using a set of tweezers for electrode attachment.

#### Electrode fabrication

To fabricate the Ag/AgCl electrodes (for ECG and CSI pseudo-reference), we screen-printed Ag/AgCl ink (91640434 - SunChemical) onto PI tape (5413 – 3M) using a rectangular-patterned silk screen (48 rectangles, each 6 x 30mm in size). To create a conductive working electrode, we screen-printed carbon ink (91592299 - SunChemical) onto PI tape (5413 – 3M) using a rectangular-patterned silk screen (48 rectangles, each 6 x 30mm in size). We left the inks to dry in a ventilated area overnight. For pH sensitivity, we deposited PANI onto the carbon through potentiostatic electrodeposition. We submerged each rectangular carbon electrode in a solution of 0.1 M aniline and 0.3 M oxalic acid. We attached the carbon electrode to the cathode of a signal generator and a stainless-steel foil electrode to the anode. We applied a potential of 1 V for 100 minutes. After polymerization, we left the electrode to dry to maximize adhesion, after which we gently rinsed it with DI water to remove any excess aniline on the polymer surface. We then left the PANI-coated carbon to condition in PBS overnight. For Li^+^ sensitivity, we prepared a Li^+^ membrane cocktail by mixing 330 mg of PVC with 660 mg of ortho-Nitrophenyloctylether (o-NPOE), 10 mg of Potassium Tetrakis (4-chlorophenyl) borate, and 5.7 mg of Li^+^ Ionophore VI in 3 ml THF. After stirring overnight, we added 20 μL of 1 M LiCl to improve sensor stability. We screen-printed the Li^+^ cocktail onto the carbon using a rectangular mask (5 x 31mm) cut into a plastic sheet (Polyethylene Terephthalate, 0.1 mm, film – Merck) using a cutter plotter. We blocked a small portion of the carbon electrode from the mask to allow for electrical connection to the circuit board (PVC insulates the carbon layer). Once screen-printing was complete, we left the electrode to dry in a ventilated area overnight. We mechanically compressed the electrode for one hour to improve physical contact between the membrane and carbon film. We then placed the electrode in DI water overnight to wash away any contaminants and unbound Li^+^ in the membrane, after which it was left to dry. All electrodes were cut into their final electrode dimensions using a cutter plotter.

### Electrode attachment

We positioned the electrodes such that the top border of the electrodes overlapped with the substrate connection point. We deposited Ag/AgCl ink on the electrode and substrate connection point to create a conductive bridge and left the ink to dry in a ventilated area. We then deposited flexible epoxy on top of the Ag/AgCl bridge and left to cure overnight.

### Characterization experiments

#### Substrate hysteresis experiment

To characterize the elastic properties of both PI substrate materials, we connected each sample to a programmable tensile and compression tester (MultiTest 5-xt - MecMesin). We taped each end of the sample to a force plate and positioned the top plate to vertically align the sample between plates. From here, we executed an automated program: the displacement of the top plate decreased to 1 mm from the bottom plate, effectively folding the sample in half. The plate remained here for 60 seconds, after which it returned to its original position.

#### Electrical characterization

We measured the frequency behavior of the antenna (S11 trace, resonant frequency, bandwidth, Q factor) using a vector network analyzer (NanoVNA) connected to an inductive antenna probe. The bandwidth is defined as the difference in frequencies 3 dB below the resonant frequency peak. The Q factor is defined as the resonant frequency divided by the bandwidth. To measure the magnitude of power induced by the designed antenna, we connected a 100 Ω resistor in parallel to the antenna and measured the voltage across the resistor at various fixed distances from an NFC reader (Adafruit PN532 breakout board). We then converted the voltage to current using Ohm’s law. To measure the power consumption of the MiFi, we connected a 1 Ω shunt resistor between the circuit and GND pin of the NTAG I2C plus. We powered the MiFi by placing it on the NFC reader, then measured the voltage across the shunt resistor and converted it to current (equivalent to load current). We measured the reading distance by placing the MiFi at increasing fixed distances from the NFC reader and checking for stable operation (1000 continuous samples) until communication ceased. We positioned the MiFi at fixed distances by adhering it to the top plate of a programmable tensile and compression tester and defined an automated program to reproducibly change the distance at fixed intervals.

#### Characterization of pH sensor

We first characterized the electrodes using a commercial potentiostat (Palmsens4). For the working electrode (PANI-C), we placed the electrodes in a beaker filled with 20 ml of PBS and left them to condition overnight. The following day, we connected the electrodes to the potentiostat and placed them on a magnetic plate to enable continuous stirring of the solution. We took OCP measurements relative to a standard double-junction Ag/AgCl electrode, during which we changed the pH using 0.1 M sulfuric acid (H2SO4) to decrease the pH and 0.1 M sodium hydroxide (NaOH) to increase pH. We simultaneously recorded the pH of the solution using a commercial pH probe (HI-5521 – Hanna instruments).

We calculated a calibration curve by plotting the potential of the electrodes against each measured pH and finding the line of best fit. The slope of the line of best fit is equivalent to the sensitivity of the electrodes. We then averaged the slope of each electrode to find the mean pH sensitivity. To characterize electrode stability (for working and reference), we connected the electrodes to the potentiostat and placed them in a beaker filled with 20 ml of PBS. The potentiostat recorded the OCP of the electrodes for at least 24 hours. We calculated the average drift of the electrodes by fitting a line to recorded data, calculating the slope of the line, and taking the average of the slopes. When integrated into the MiFi, the same protocol was followed to calculate sensor stability and sensitivity. Instead of using a commercial potentiostat, however, we placed the MiFi at the bottom of the beaker. We also placed an NFC reader directly underneath the beaker to enable powering and data communication/recording. We used a one-point calibration (using the potential in 1x PBS: ∼7.4 pH) to align the MiFi and potentiostat calibration curves for comparison.

#### Characterization of Li^+^ sensor

We characterized the Li^+^ sensor following the same protocol as the pH sensor. We conditioned the working electrodes in a beaker filled with 20 ml of PBS (+ 1mM LiCl) overnight. We obtained the calibration curve by adding increasing volumes of 1 M LiCl to spike the Li^+^ concentration of the solution at known concentrations. To characterize electrode stability, we placed the electrodes in a beaker filled with 20 ml of PBS spiked with 1mM LiCl. When integrated into the MiFi (Config 3), we followed the same protocol to calculate sensor stability and sensitivity. We used a one-point calibration (using the potential at 1mM LiCl) to align the MiFi and potentiostat calibration curves for comparison.

#### Characterization of ECG monitor

We connected two leads of a signal generator into either side of a sample of chicken tissue, roughly five centimeters apart. We set the input signal as a simulated ECG signal with 30mVpp amplitude and frequencies ranging between 6-8 Hz. We placed the MiFi (Config 1) on the chicken tissue between the signal leads to detect the transmitted signal. We then placed an NFC reader underneath the chicken tissue to enable powering and communication/recording.

### Multiplexed in vitro experiments

All configurations of the MiFi followed the same experimental setup (see SI for illustration): we placed the device at the bottom of a beaker filled with 20 ml of PBS and the beaker was placed on a magnetic plate to enable continuous stirring. We placed an NFC reader underneath the beaker to enable powering and communication. During recording, we induced a change in each sensing module consecutively to enable monitoring of cross-interference. For all non-CSI modules, we monitored for approximately 10 minutes after each stimulus. We extended the time for CSI stimuli to allow the chemical addition to mix homogenously within the solution. For the temperature sensor, we increased the temperature of the solution by turning on a blow dryer for 5 seconds approximately 5 cm away from the beaker. We decreased the temperature of the solution by placing a cold damp towel around the beaker until the temperature drops by approximately 1 °C. For the ECG monitor, we applied a potential signal across the solution by submerging aluminum foil electrodes at either side of the beaker and connecting them to a signal generator. We transmitted a 1Vpp sine wave at 2, 4, and 6 Hz.

For the accelerometer, we taped a vibration motor (JYC1434) and driver (DRV2605 - TI) underneath the NFC reader antenna to create an inertial stimulus. We configured the driver to spike at 100% power at 2, 4, and 6 Hz. For the pH sensor, we increased the pH of the solution by spiking the solution with 0.2 M disodium phosphate (Na2HPO4). We decreased the pH of the solution by spiking the solution with 0.1 M citric acid. For the Li^+^ sensor, we increased the Li^+^ concentration of the solution by spiking the solution with 1 M LiCl. We decreased the Li^+^ concentration by diluting the solution with PBS.

### In vivo experiments

All procedures were carried out under the terms and conditions of licenses issued by the UK Home Office under the Animals (Scientific Procedures) Act 1986. All in vivo experiments were performed using Sprague-Dawley rats. Rats were anesthetized with urethane (1.35g/kg, IP), using isoflurane for induction. Rats were placed on a small animal physiological monitoring system (Harvard Apparatus) during the operation to enable commercial monitoring of the animal’s physiology. The internal body temperature of the animal was also maintained using the monitoring platform. After induction, atropine (0.66 ml/kg, 1% w/v) was administered subcutaneously to reduce mucous secretions. Animals were humanely killed in accordance with Schedule 1 to the Animals (Scientific Procedures) Act 1986.

### Config 1 experiment

We subcutaneously implanted Config 1 of the MiFi directly above the heart to capture the depolarization wave of the heartbeat. The data were collected while the animal was kept at rest in an anaesthetized state. To extract the cardiac waveform from the data, we took the FFT and isolated the frequency peaks physiologically related to the ECG waveform (for example, excluding peaks related to breathing and musculoskeletal noise – see SI for FFT visualization). We then reconstructed the cleaned signal in the time domain using the inverse FFT. To compare the waveform of each heartbeat, we plotted the average and standard deviation of the waveforms extracted from the MiFi data against the average of the ECG waveforms collected from the commercial monitoring system during the same period. To calculate the pH of the ISF, we performed a one-point calibration using the potential at a known pH, after which we converted the potential data to pH using our derived calibration curve. To remove high frequency noise, we applied a moving average filter with a window length of 10 seconds.

### Config 2 experiment

The MiFi was implanted subcutaneously under the left forelimb where sensitivity to heart and breathing motion was found to be maximized (see SI for further details). After keeping the animal at rest in an anaesthetized state for approximately 10 minutes, 2% isoflurane was administered by mask delivery and maintained for a further 15 minutes.

Following this, the isoflurane supply was removed and the animal was left in this state (under urethane anesthesia) for a further 15 minutes. The animal was culled using an IP injection of pentobarbital (2 ml). To calculate the pH of the ISF, we used the same protocol as defined for Config 1. For the accelerometer data, all data processing was done using a moving window of length 1 minute and 55 second overlap. To extract the breathing pattern, we applied a moving bandpass filter (passband = 0.5-2 Hz). The breathing rate was estimated by finding the period between peaks of the breathing pattern, converting to bpm (breaths per minute) to get the instantaneous breathing rate, then averaging these values to reduce the influence of noise. To extract the cardiac pattern, we first applied a moving high pass filter (fc = 4 Hz) to remove the breathing artifacts from the signal. Then, we took the FFT of the data, isolated the frequency peaks related to the motion of the heart, and reconstructed the signal using only these peaks (see SI: Fig S12). To increase the accuracy of heart rate detection, we increased the prominence of the S1 peaks by scaling the FFT-cleaned signal (scaling factor = 10*mean(window)^0.25^). The heart rate was calculated by finding the period between S1 peaks, converting to bpm (beats per minute) to get the instantaneous heart rate, then averaging these values to reduce the influence of noise.

For more effective comparison between the MiFi and commercial monitoring system, we applied a low pass filter (fc = 2 Hz) to the commercial breathing signal and a moving average filter (window = 1 minute) to the commercial heart rate signal.

### Config 3 experiment

The MiFi was implanted subcutaneously at the scruff. The data was collected while the animal was kept at rest in an anaesthetized state. We let the sensor condition to its environment for 30 minutes before recording, after which we recorded for 40 minutes without intervention to establish a baseline. We then injected 100 mg/kg LiCl (rat weight: 719 g) through IP injection. To calculate the Li^+^ concentration in the ISF, we first applied a moving median filter (window = 10 seconds) to remove high frequency noise. To remove systemic non-physiological artefacts from the Li^+^ data, we isolated the artefacts using the pH data as a reference and subtracted them from the Li^+^ data. To remove quantization noise from ADC sampling, we also applied a moving average filter (window = 5 minutes). We then converted the potential data to Li^+^ concentration using our derived calibration curve.

## Acknowledgements

F.G. would like to thank Imperial College London, Department of Bioengineering and Imperial College Centre for Processable Electronics. S.O. acknowledges the Imperial President’s PhD Scholarship. F.G. and P.C. thank EPSRC Centre for Doctoral Training in Plastic Electronics (EP/l016702/1). F.G. also acknowledges support from UKRI Cure4Aqua (Ref: 10050496). A.S. and A.S.C. would like to thank Fu Siong Ng for the use of his laboratory and the British Heart Foundation (WHCF-PSJ123) to carry out *ex vivo* validation tests.

## Author Contributions

S.O. and F.G. conceived the structure of the manuscript. S.O. led the writing of the manuscript and experimental work. J.D.G., A.N., P.C., I.C.C., F.A., A.S., M.M., A.S.K., V.S., and A.S.C. contributed to experimental work. J.D.G., A.N., E.A., and A.S.K. contributed to the writing of the manuscript. All authors reviewed and agreed on the manuscript before submission.

## Data Availability Statement

The data that support the findings of this study are available from the corresponding author upon reasonable request.

