## Supplementary Information for "A Geometrically Transient Material for Bioelectronic Implants"

Supplementary Figures and Legends S1 – S16	6

References	14

### 1. Details of circuit layout for foldable circuit board

We optimized the locations of the ICs to minimize the number of columns with (relatively) tall components. Analyzing the densest circuit configuration (Config 1), we organized four out of five ICs into one column: the microcontroller was only left in a separate column due to connection layout requirements. To prevent two ICs from folding directly on top of one another – which would increase the width of the device by appx. 0.5mm per fold (excluding encapsulation) and place additional stress on the components during folding – we created a space along the midpoint of the column for the microcontroller to occupy during folding. With this design, no additional stresses were propagated and the height of the microcontroller did not contribute to the overall thickness of the folded device. The rest of the columns defined had either 0201 components only (< 0.3mm in height) or no components at all (just antenna), so have a negligible impact on the final width of the device. To minimize device thickness when folded, we only encapsulated folding columns populated with components (four out of six).

### 2. Programming Arduino core and firmware for MiFi platform

The ATtiny20 does not come with its own dedicated firmware and library packages that allow it to be easily integrated into popular prototyping environments such as Arduino, PlatformIO, etc. Thus, we constructed a custom Arduino core for the ATTiy20 based off of the open-source cores available for other ATTiny models, specifically the ATtiny10Core from technoblogy^1^. In addition to building the Arduino core, we developed a custom programming tool to enable on-board programming and to keep the microcontroller re-programmable at most stages of fabrication (i.e. until encapsulation). The ATtiny20 contains 2kB of in-system re-programmable flash that can be written to use the Tiny Programming Interface (TPI). In addition to VCC and GND, TPI programming is executed using a three-pin interface: TPICLK (clock input), TPIDATA (data input), and RESET (enables programming). We purchased and modified a USBASP AVR Programmer and adapter (Hobby Components) to create a three-pin interface. To do this, the MOSI and MISO pins were connected to the same bus: each of these pins support unidirectional communication but the TPIDATA pin supports bi-directional communication. All other connections were kept the same. Finally, to interface the programmer with the circuit board, we connected spring-loaded male header pins to the adapter, allowing for continuity between the on-board connection pads and programming without any physical damage to the circuit board.

While the ATtiny20 possesses built-in hardware for serial peripheral interface (SPI) communication, there is no hardware for I2C. Software implementations of I2C (known as ‘soft I2C’) was enabled using the open-source library called SoftI2CMaster designed by felias-fogg using any two defined GPIO pins^2^. It should be noted that, similar to the hardware implementation, pull-up resistors are still required for the SDA and SCK pins to enable differentiation between ‘HIGH’ / ‘1’ and ‘LOW’ / ‘0’ states. We kept communication with all digital peripherals as simple and lightweight as possible by directly addressing the specific registers needed for configuration and data reading/writing rather than relying on pre-developed libraries. To sample the analog sensors, we used the built-in 8-bit and 16-bit timers to create interrupt timers that could sample the sensors at precisely defined periods. Lower sampling frequencies, such as those used for pH and ion concentration (< 10 Hz) require the 16-bit timer as the counter must wait more than 255 ticks (maximum number stored in 8-bit value) of the CPU’s clock before it can reset.

To improve data sampling rates, specifically for ECG, we modified the NFC configuration of the NTAG I2C PLUS to transmit data from the microcontroller directly to the SRAM for wireless transmission. To reach the achieved data rates, we reduced the size of the data packet to 32 bytes instead of filling the full 64-byte SRAM buffer. Since multiple sensors were sampled within one device, each 32-byte data package was structured using a fixed template prior to transmission. This helped the NFC-reader easily identify which data belongs to which sensor. Additionally, although NDEF is typically used to transmit data, the overhead required for formatting takes up too much of the NTAG’s limited SRAM, so each sensor value was transmitted as two 8-bit values and converted to decimal values by the NFC-reader.

### 3. CSI circuit design

We used a dual op-amp IC (ADA4505 – Analog Devices: 1.42 x 1.42 mm) to configure two buffers, used for Vref and the OCP output. The buffer electrically isolates the electrode from the rest of the circuit and prevents the low impedance of the ADC from loading and consequently attenuating the signal. From there, the signal is digitized by one of the 10-bit ADC channels of the ATtiny20. Due to the stability and relatively large sensitivity of potentiometric sensors (~59 mV/pH as predicted by Nernst equation) relative to the precision of the ATtiny20’s ADC (3 mV/step on average), no additional pre-processing or amplification steps were added to the analog front end of the sensor. Vref is created using a voltage divider between VCC and GND and is set to half of VCC (~1.45 V).

### 4. ECG circuit design

The high-pass filter is pulled up to Vbuf to access the full amplification range of the IA. A 2.2nF capacitor is placed in parallel to the electrodes to help suppress high-frequency EMF noise. Similar to the CSI, the output of the IA is biased to Vref to capture the amplified positive and negative changes in signal. The final amplified ECG signal is connected to one of the ATTiny20’s 10-bit ADC channels for digitization. A 1 µF capacitor is placed between the supply voltage and ground rails to prevent browning out during NFC operation and modulation: browning out resets the registers of the NFC chip and removes the high-speed transmission settings required to send data sampled at high frequencies, such as ECG. An additional 0.1 µF capacitor is used to suppress high frequency noise.

**Table S1.** **Price breakdown of MiFi fabrication at two scales: one device, x1000 devices (or closest available price)**

| **Materials** | **Amount/Size** | **Buying x1 (USD)** | **Buying x1000 (or closest to, USD)** |
| --- | --- | --- | --- |
| **Circuit board** | | | |
| Polyimide tape | 400mm^2 | 0.162 | 0.162 |
| Polyimide sheet | 400mm^2 | 0.176 | 0.176 |
| Copper foil | 166.8mm^2 | 0.014 | 0.014 |
| Solder (260C) | ~0.05g | 0.078 | 0.078 |
| Solder (200C) | ~0.01g | 0.015 | 0.012 |
| **TOTAL** |  | **0.501** | **0.498** |
| **IC components** | | | |
| Attiny20 | 1 | 1.134 | 1.053 |
| ADA4505-2 | 1 | 1.809 | 1.102 |
| NTAG I2C PLUS | 1 | 1.445 | 0.626 |
| Config 1 | | | |
| AS6212 | 1 | 1.863 | 0.764 |
| MAX41400 | 1 | 2.862 | 2.066 |
| Config 2 | | | |
| AS6212 | 1 | 1.863 | 0.764 |
| BMA530 | 1 | 2.862 | 2.0655 |
| **TOTAL: Config 1** |  | **9.113** | **5.611** |
| **TOTAL: Config 2** |  | **7.790** | **4.439** |
| **TOTAL: Config 3** |  | **4.388** | **2.781** |
| **Passive components** | | | |
| 100pF | 1 | 0.378 | 0.090 |
| 0.1uF | 1 | 0.108 | 0.015 |
| 1uF | 1 | 0.108 | 0.012 |
| 4.7k | 2 | 0.475 | 0.073 |
| 1M | 2 | 0.410 | 0.059 |
| Config 1 | | | |
| 0.1uF | 1 | 0.108 | 0.015 |
| 1M | 2 | 0.410 | 0.059 |
| 0.33uF | 2 | 0.454 | 0.097 |
| 2200pF | 1 | 0.378 | 0.084 |
| **TOTAL: Config 1** |  | **2.016** | **0.363** |
| **TOTAL: Config 2** |  | **1.096** | **0.185** |
| **TOTAL: Config 3** |  | **1.096** | **0.185** |
| **Encapsulation** | | | |
| Resin | ~0.1 g | 0.003 | 0.003 |
| Parylene-C | negligible | negligible | negligible |
| **TOTAL** |  | **0.003** | **0.003** |
| **Electrodes** | | | |
| Polyimide tape | 32.4 mm^2 | 0.003 | 0.003 |
| Config 1 | | | |
| Ag/AgCl ink | 3 x 9 mm^2 | 0.022 | 0.022 |
| Carbon ink | 7.2 mm^2 | 0.003 | 0.003 |
| PANI (oxalic acid, aniline) | 7.2 mm^2 | 0.004 | 0.004 |
| Config 2 | | | |
| Ag/AgCl ink | 1 x 9 mm^2 | 0.007 | 0.007 |
| Carbon ink | 7.2 mm^2 | 0.003 | 0.003 |
| PANI (oxalic acid, aniline) | 7.2 mm^2 | 0.004 | 0.004 |
| Config 3 | | | |
| Ag/AgCl ink | 3 x 9 mm^2 | 0.007 | 0.007 |
| Carbon ink | 7.2 mm^2 | 0.003 | 0.003 |
| PANI (oxalic acid, aniline) | 7.2 mm^2 | 0.004 | 0.004 |
| Li-selective membrane (Li ionophore VI, PVC, o-NPOE, NaTPB, THF, LiCl) | 13.5 mm^2 | 0.593 | 0.593 |
| **TOTAL: Config 1** |  | **0.032** | **0.032** |
| **TOTAL: Config 2** |  | **0.017** | **0.017** |
| **TOTAL: Config 3** |  | **0.610** | **0.610** |
| **TOTAL COST: CONFIG 1** |  | **12.313** | **6.577** |
| **TOTAL COST: CONFIG 2** |  | **9.733** | **5.150** |
| **TOTAL COST: CONFIG 3** |  | **6.923** | **4.084** |

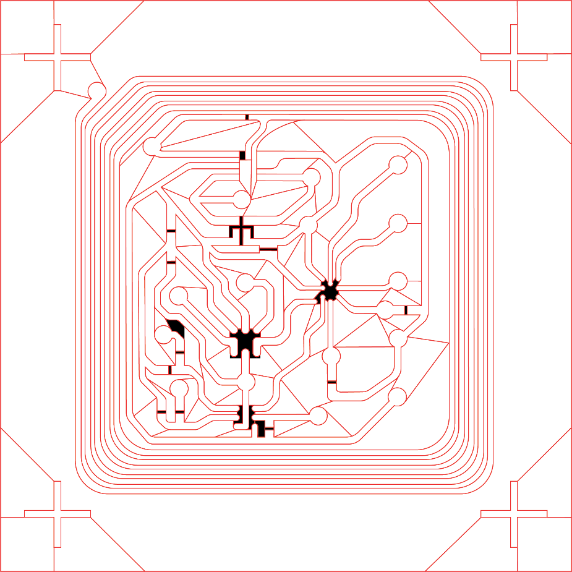

**S2.** **Design layout of engrave and cut guidelines for laser cutting the top layer of copper connections of the MiFi circuit board.** The (RGB) black filled shapes represent the areas to be engraved and the (RGB) red lines represent the lines to be cut by the laser cutter. Additional cut lines are added to aid with copper foil peeling.

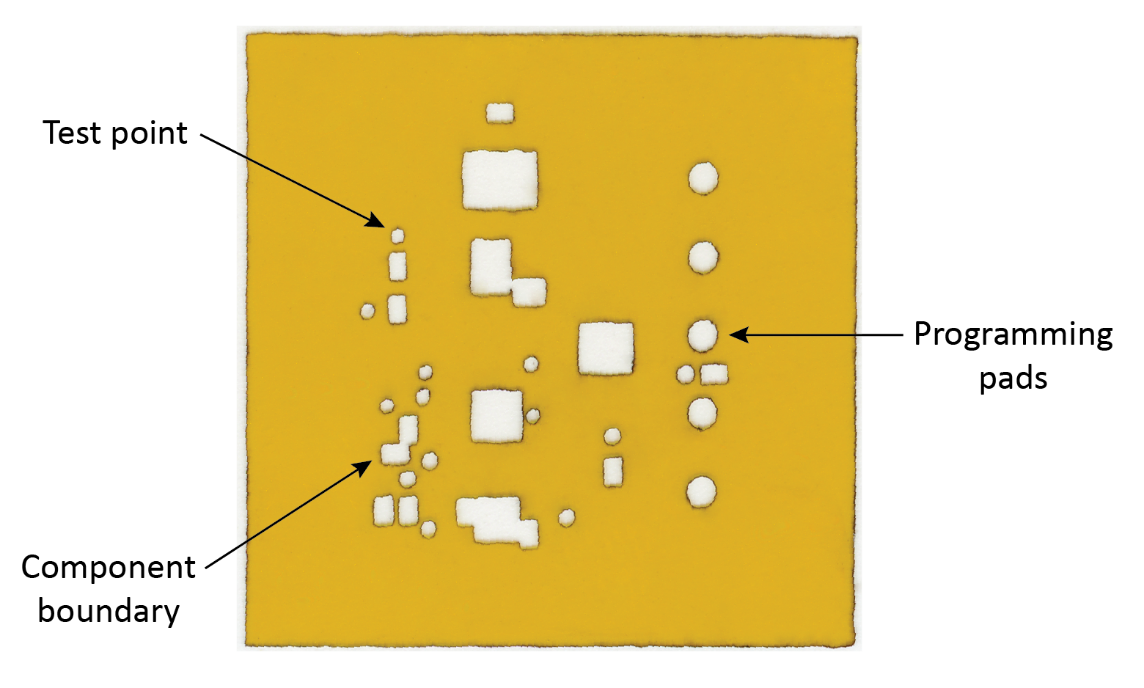

**S3. Photograph of PI template mask.** The template mask provides more precise positioning of small components (< 0.5mm) during soldering by acting as a boundary around the solder pads to avoid component movement. We laser cut small circles above copper traces to create test points for checking connections and overall circuit function. We also laser cut a column of larger circles on the right side of the circuit board to enable on-board programming of the microcontroller.

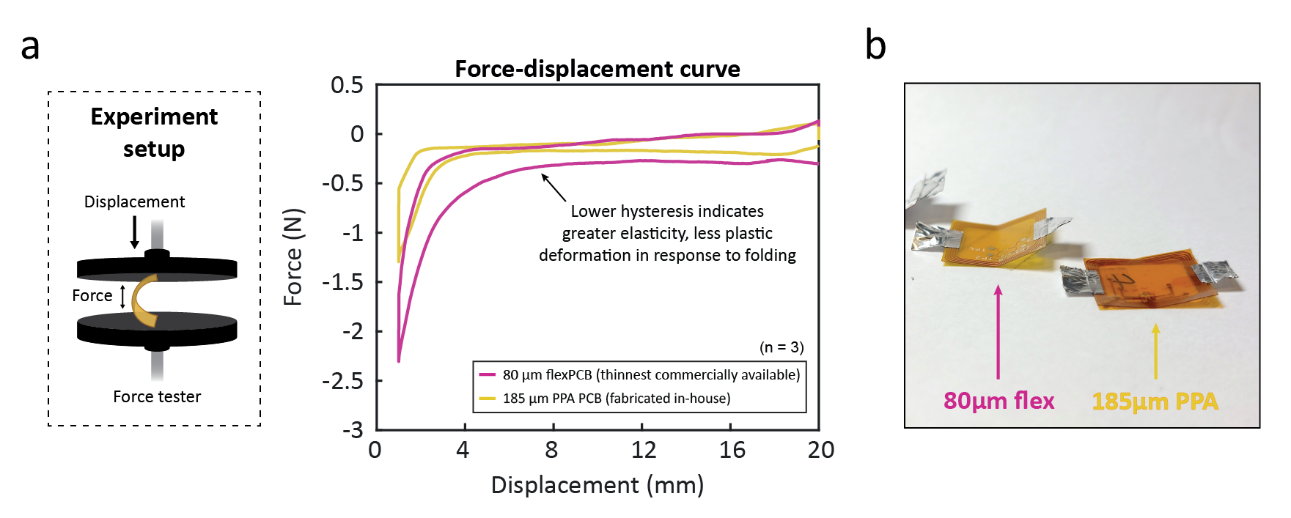

**S4. Substrate hysteresis experiment to investigate material properties of two PI substrates for folding applications.** (a) Illustration of experiment setup (left) and resulting force-displacement curve (right). Samples were folded to 1 mm bending radius using a programmable tensile testing machine for 1 minute and then left to relax for 5 minutes. (b) Photograph of two different PI samples (80 µm commercial flexPCB and 185 µm in-house fabricated PPA PCB) after completing hysteresis experiment. Metal tape is added to each side of the sample to facilitate attachment to tensile testing machine.

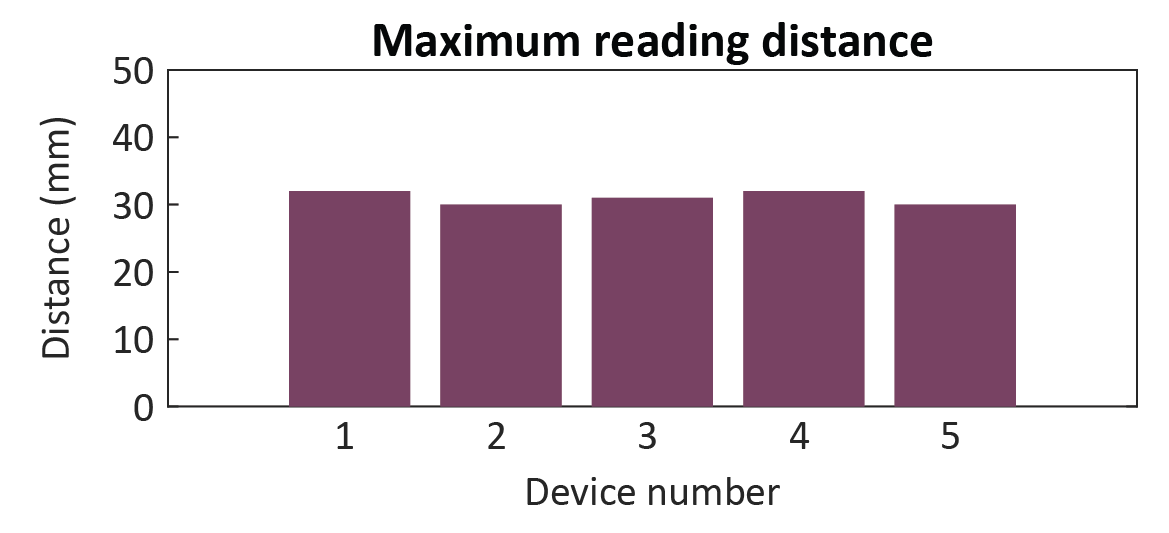

**S5. Maximum reading distance between MiFi and NFC reader before communication is lost.** 1000 samples of data must be successfully recorded and transmitted wirelessly to the NFC reader for communication to be considered successful.

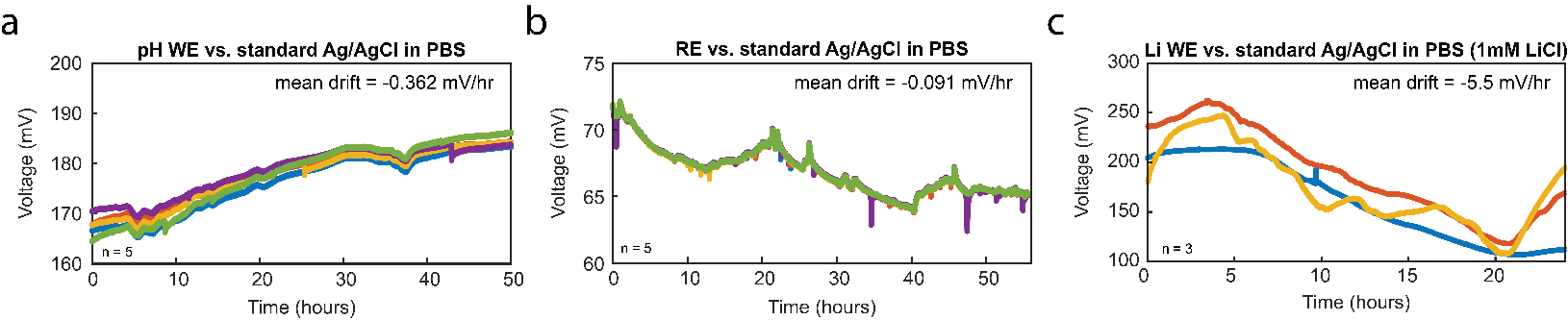

**S6.** Stability data of pH and Li working electrodes and Ag/AgCl pseudo-reference electrode against standard double-junction Ag/AgCl electrode using commercial potentiostat.

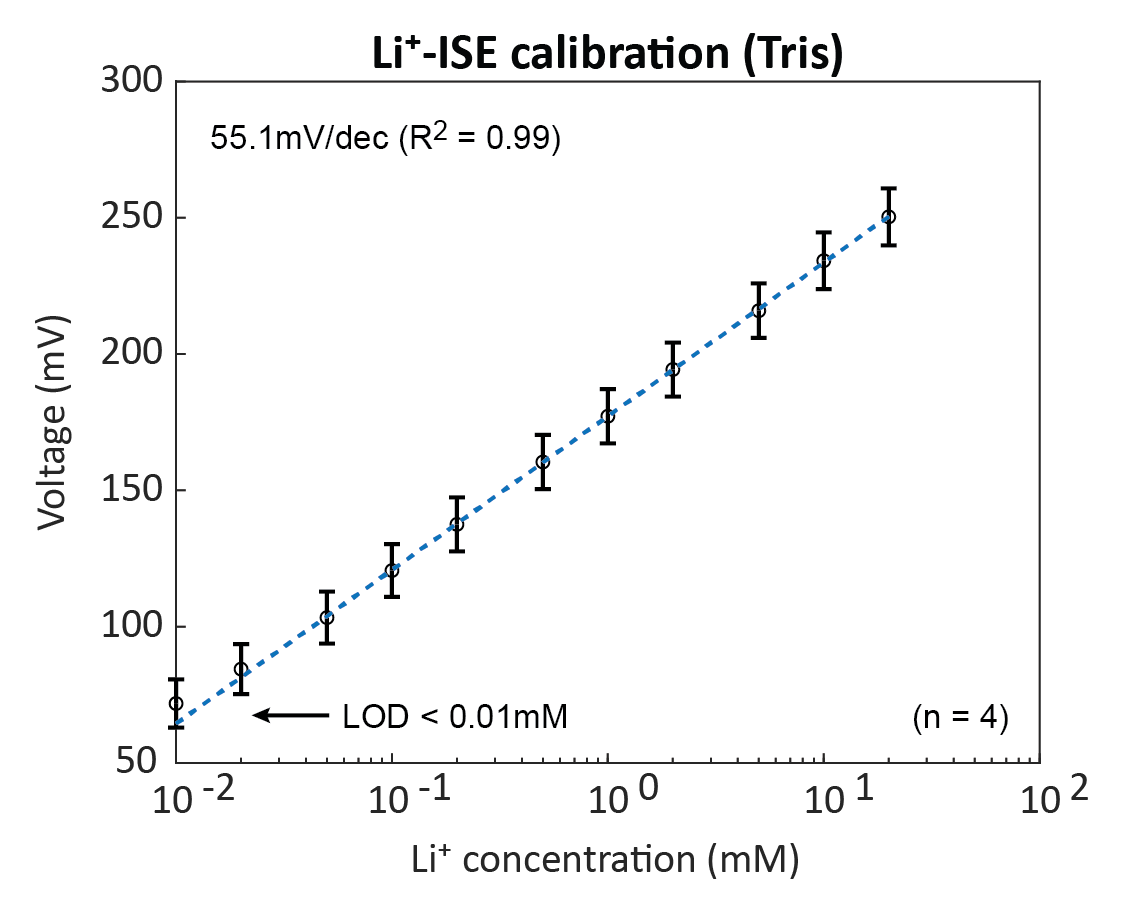

**S7. Calibration curve for Li electrochemical sensor in TRIS buffer (non-ionic solution).** We used TRIS buffer solution to characterize the electrodes in idealized conditions, where no interfering ions are present to degrade sensor performance. Electrodes demonstrated high sensitivity in response to Li addition, with values close to that predicted by the Nernst equation: 55.1mV/dec. The electrode behaviour remains within the linear region of detection for all concentrations tested, indicating a micro-molar LOD that is far below the expected serum Li levels from BPD under-dosage.

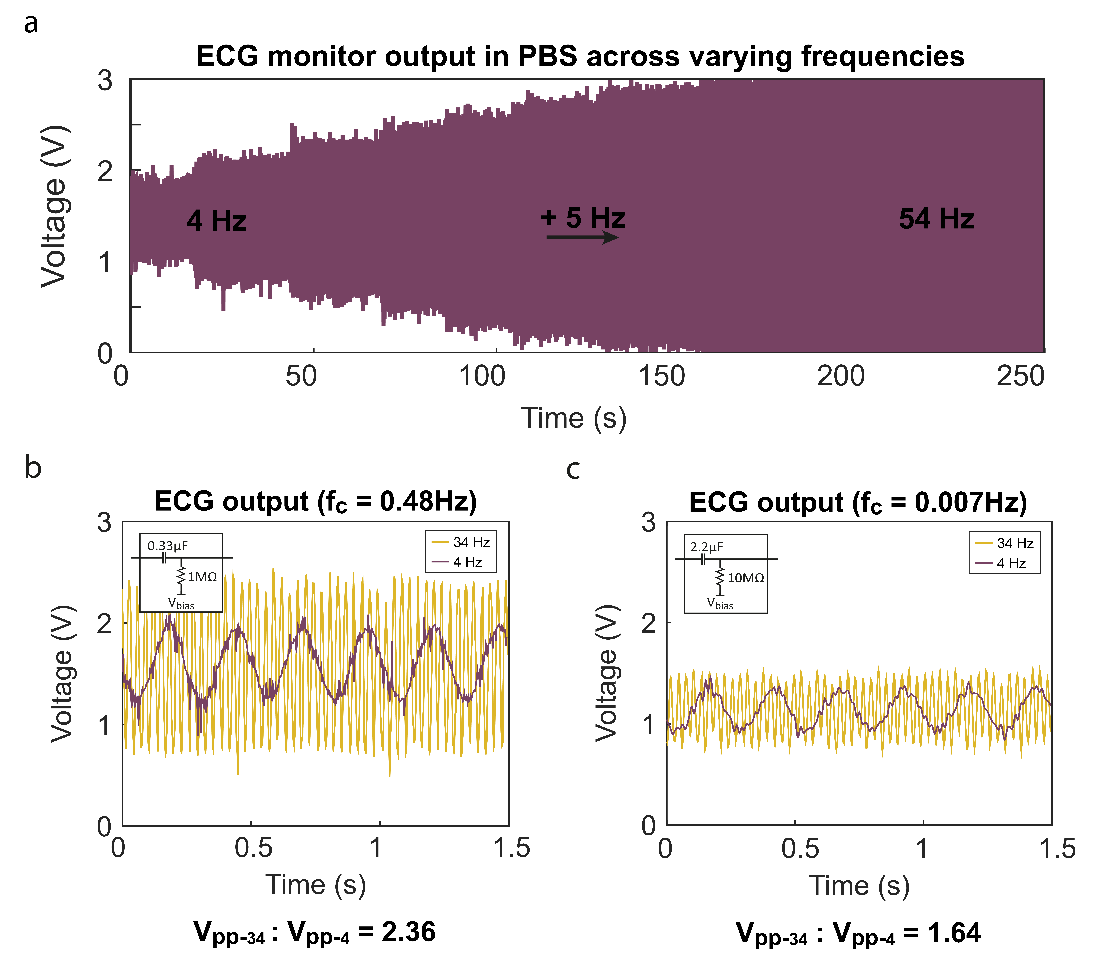

**S8. Investigation into attenuation of ECG output from high pass filter.** (a) ECG output over increasing frequency from 4 Hz to 54 Hz in 5 Hz increments (fc = 0.48 Hz, Vpp = 0.31 V). (b) ECG output at two input frequencies – 4 Hz, 34 Hz – when high pass filter set to fc = 0.48 Hz (Vpp = 0.31V). (c) ECG output at two input frequencies – 4 Hz, 34 Hz – when high pass filter set to fc = 0.007 Hz (Vpp = 0.02 V). ‘V_pp-34_ : V_pp-4_’ represents the ratio in mean amplitude between the recorded sine wave at 34 Hz compared to 4 Hz: a higher value indicates a smaller difference in amplitude so less attenuation by the filter.

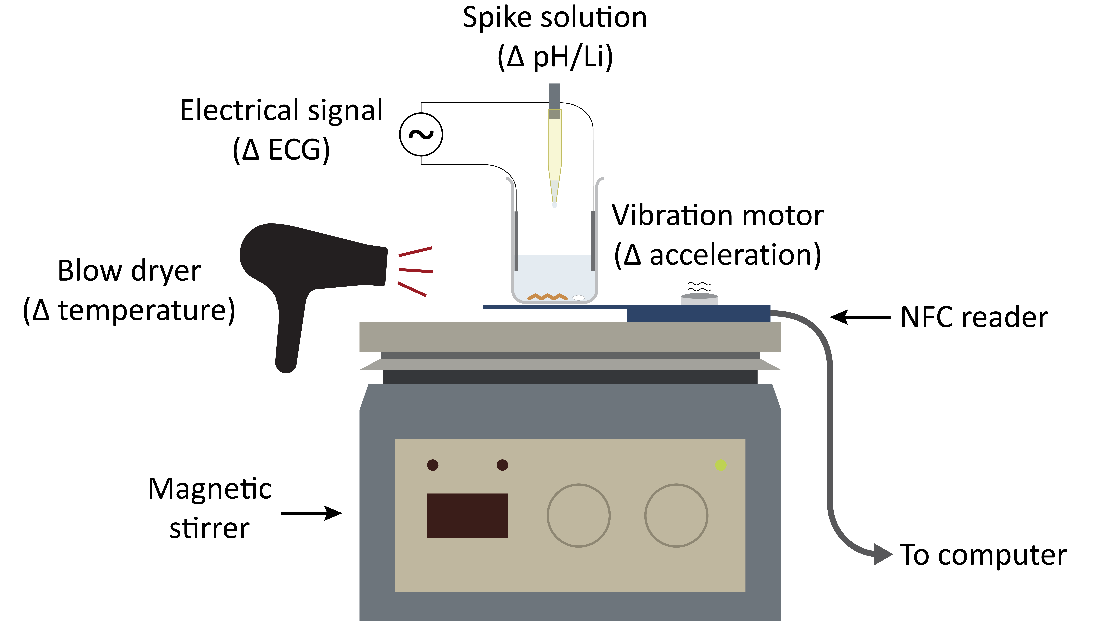

**S9. Experiment setup for in vitro multiplexing experiments of Config 1-3 of MiFi.**

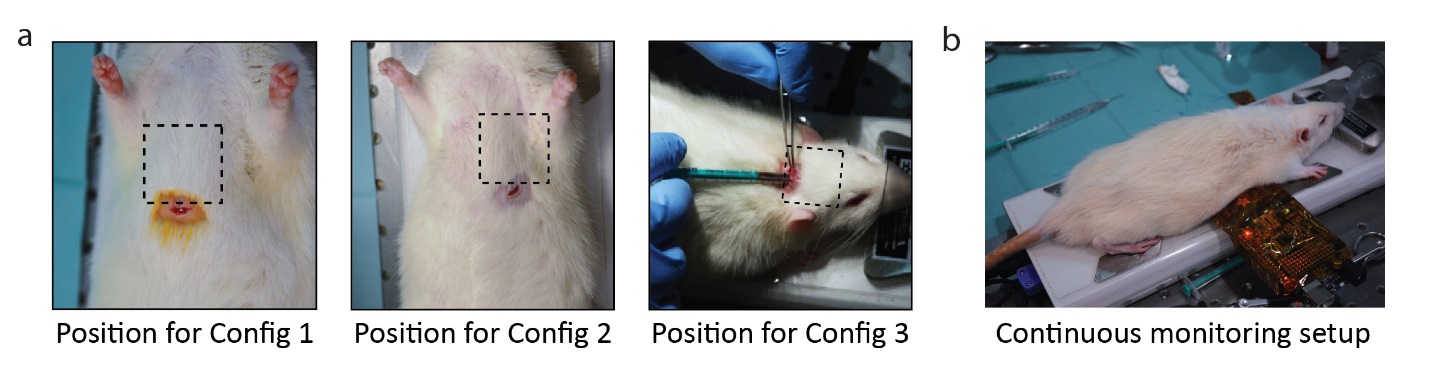

**S10. Implantation and monitoring setup for *in vivo* rat experiments.** (a) Photographs of implantation site for Config 1 (left), Config 2 (middle), and Config 3 (right) of MiFi during *in vivo* rat experiments. (b) Photograph of continuous monitoring setup. To enable simultaneous recording from the MiFi and commercial monitoring platform, the rat was positioned in the prone position so that the paws contacted the metal pads of the commercial system (necessary for ECG monitoring). An NFC reader was placed between the rat and commercial system to enable powering and wireless communication with the implanted MiFi.

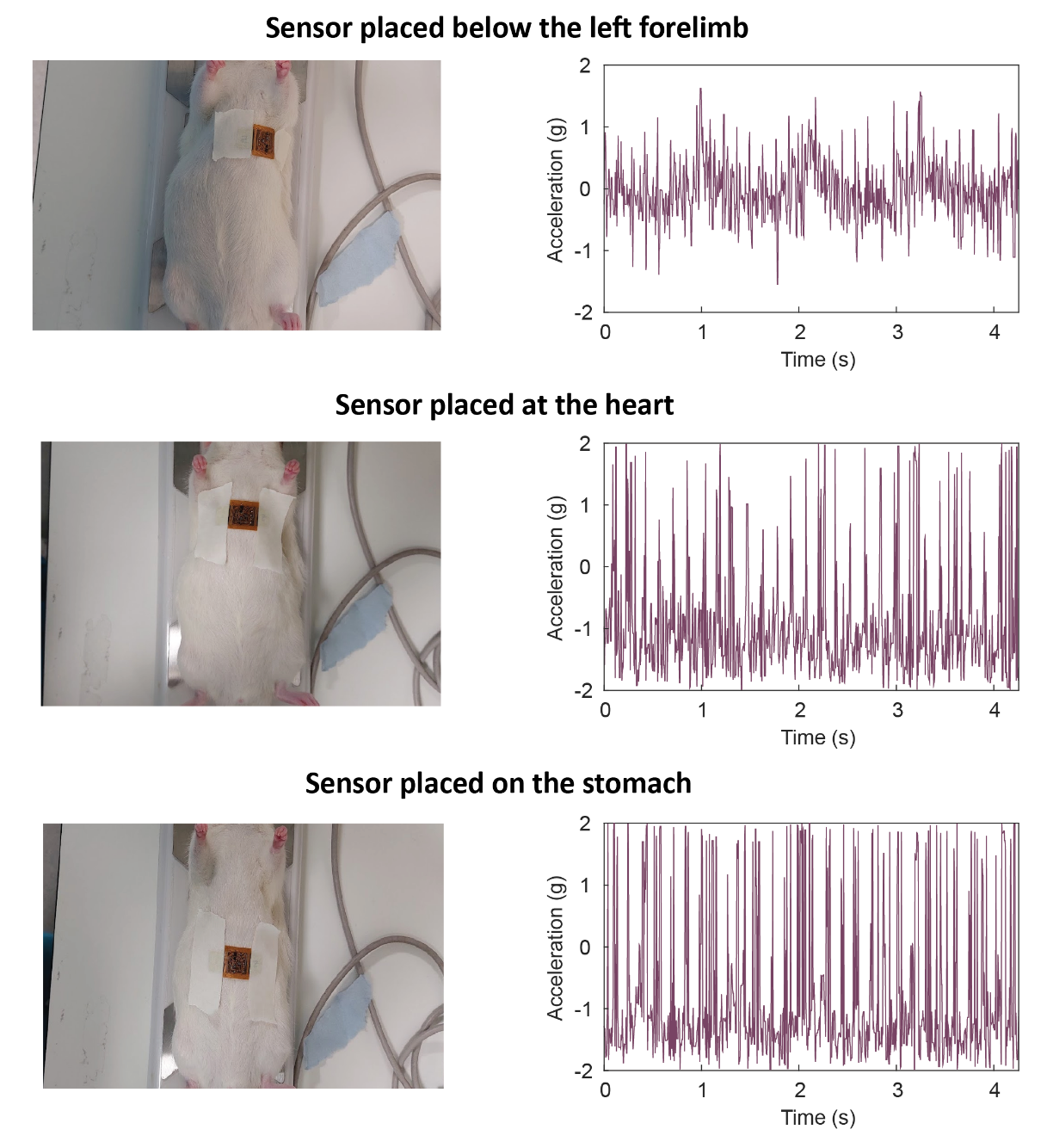

**S11.** **Noninvasive investigation into Config. 2 placement for optimal heart rate and breathing rate detection**. Left panel: photographs of device position. Right panel: accelerometer output at designated location. Three locations were tested as potential implantation sites for heart rate and breathing rate monitoring: the location above the heart, underneath the left forelimb, and on top of the stomach. We chose the location above the heart because of the sensor’s proximity to both the heart and lungs. We chose underneath the forelimb because this location provides the best conformal fit between the body and device. We additionally investigated on top of the stomach as we believed that this location may be able to pick up the heart and breathing rate with less interference from surrounding skeletal muscle (not as dense as in the chest). A noninvasive test was carried using a Sprague-Dawley rat anaesthetized using isoflurane. The device was taped onto each of the chosen locations using a medical tape to minimize the impact of relative motion between the body and the device and to minimize noise caused by fur movement. Recording at the heart captured strong mechanical movements from the heart, but was also prone to picking up large noise artefacts that made breathing harder to distinguish and would make the heart pattern harder to isolate during post-processing. Contrary to our assumption, recording at the belly was the most noise-prone, however, it was still able to pick up both the heart and breathing rate despite its distance from the organs. Recording underneath the forelimb ultimately provided the highest clarity in signal. Although the heart patterns are less pronounced than if recording at the heart, the improved signal clarity makes it much easier to isolate. The breathing pattern underneath the forelimb is also the most pronounced. For these reasons, it was decided that proceeding *in vivo* experiments would use under the left forelimb as the implantation site.

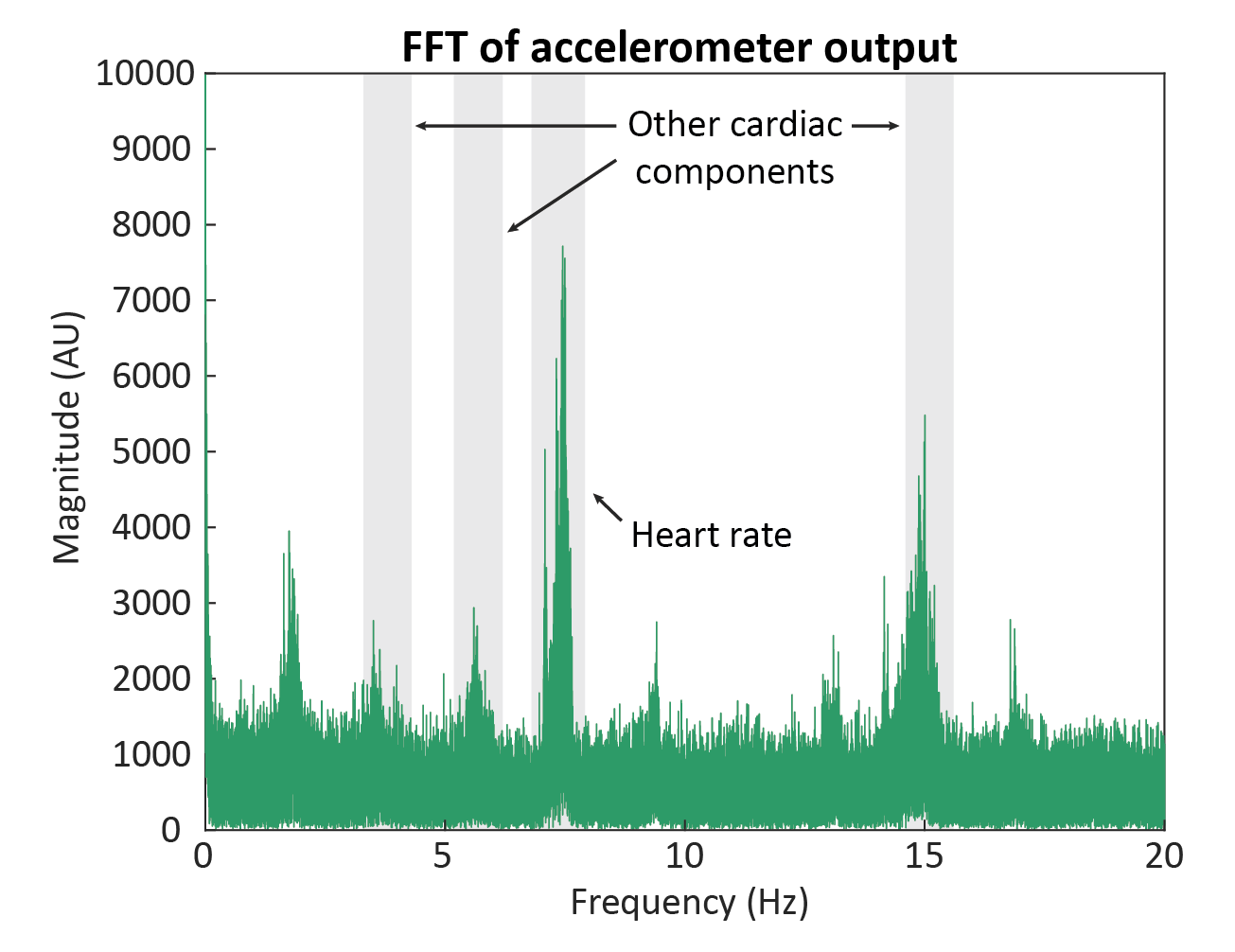

**S12.** **FFT of accelerometer data collected from Config 2 of MiFi.** Four key frequency peaks were isolated and used to reconstruct the heartbeat signal via inverse FFT: the fundamental heart rate peak (~7-8 Hz), the first harmonic peak (~15-16 Hz), and two sub-fundamental peaks (~3-4 Hz, ~ 5-6 Hz).

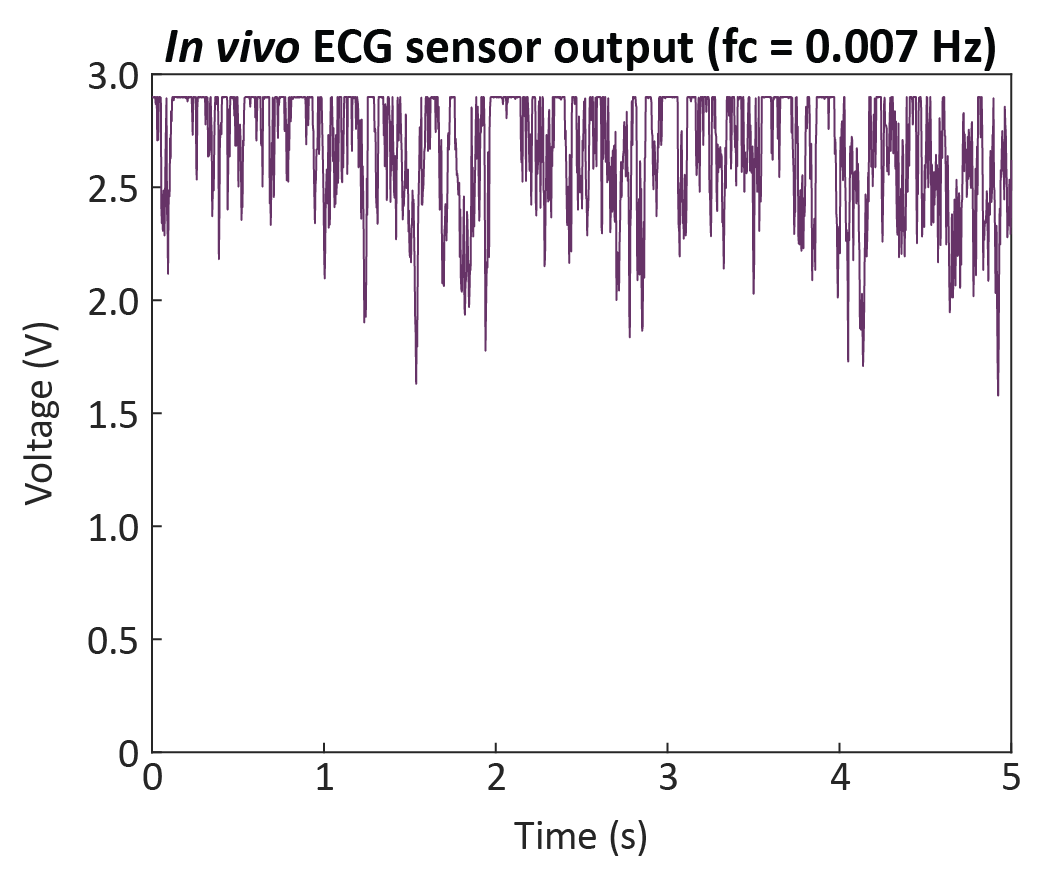

**S13. In vivo data from ECG sensor of Config 1 when high-pass filter is configured to fc = 0.007 Hz.** The cut off frequency (fc) is too low to attenuate low frequency drift between ECG electrodes, resulting in an amplified (x200) offset at the sensor output that saturates the ECG signal to VCC (~2.9V).

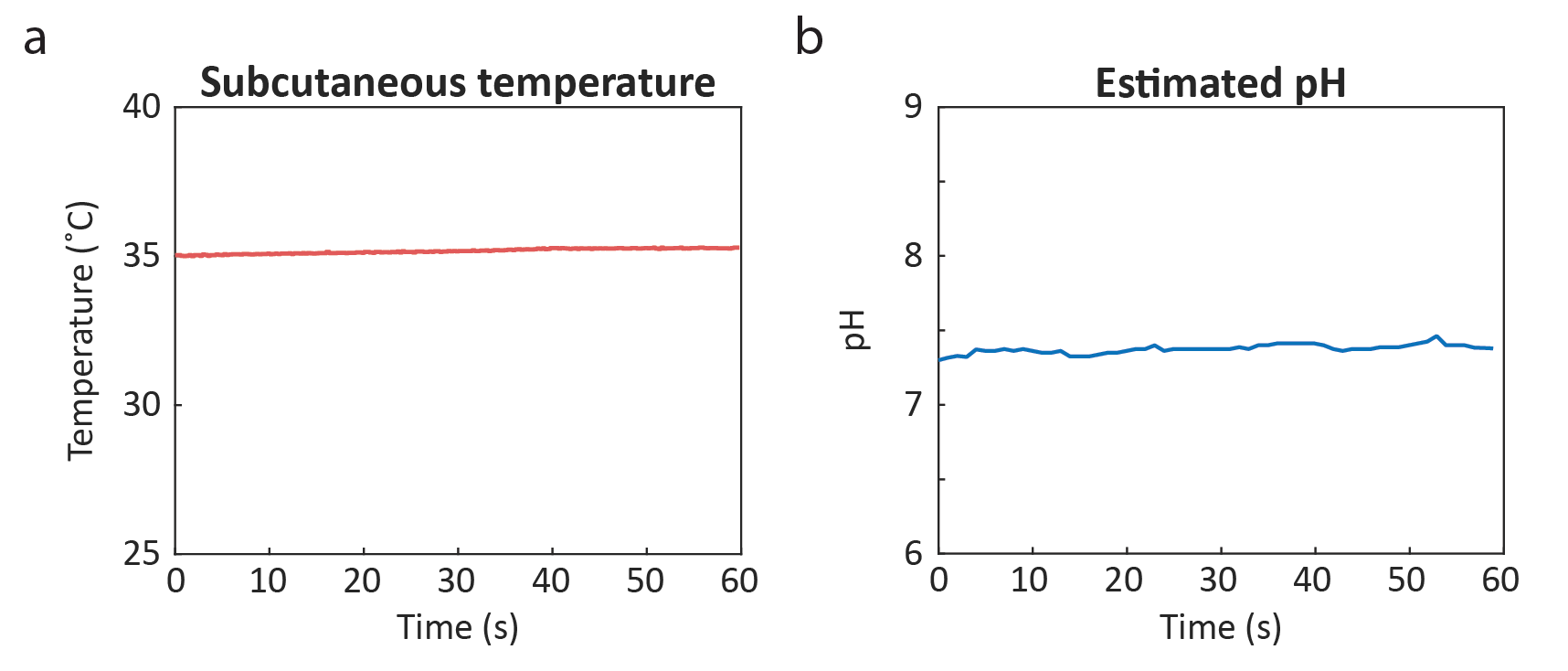

**S14.** **Remaining data collected from Config 1 of MiFi during *in vivo* experiment.** (a) Subcutaneous temperature. (b) Estimated pH of interstitial fluid.

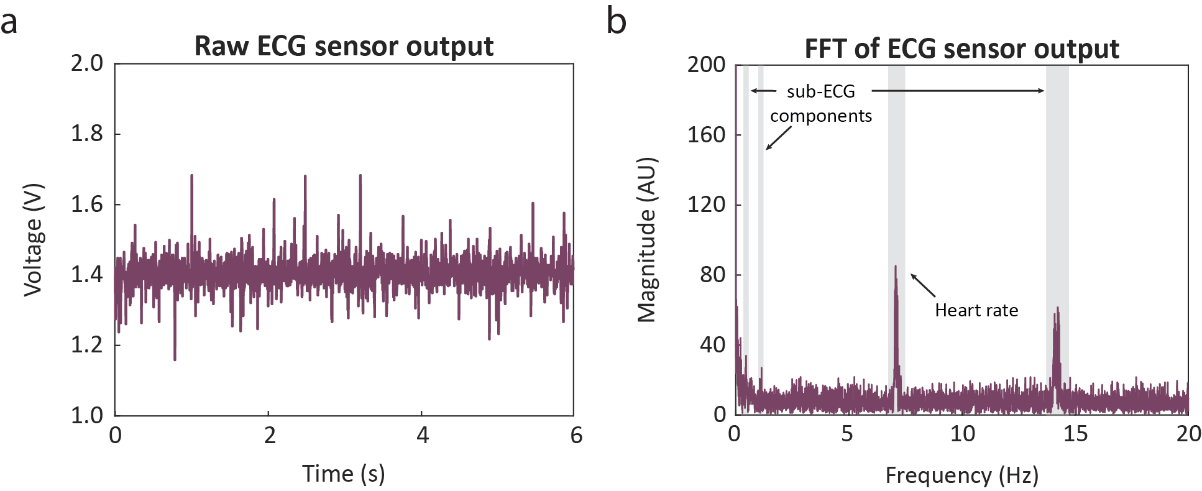

**S15.** **ECG data and analysis collected from Config 1 of MiFi during *in vivo* experiment.** (a) Clip of raw ECG data; b) FFT of ECG data, identifying the prominent peaks related to the animal’s ECG waveform that were used to reconstruct the cleaned ECG signal via inverse FFT: the fundamental frequency peak (~6.8-7.4 Hz), two harmonic peaks (~13.7-14.7 Hz, ~20.8-21.9 Hz), and one sub-fundamental peak (~0.35-0.85 Hz).

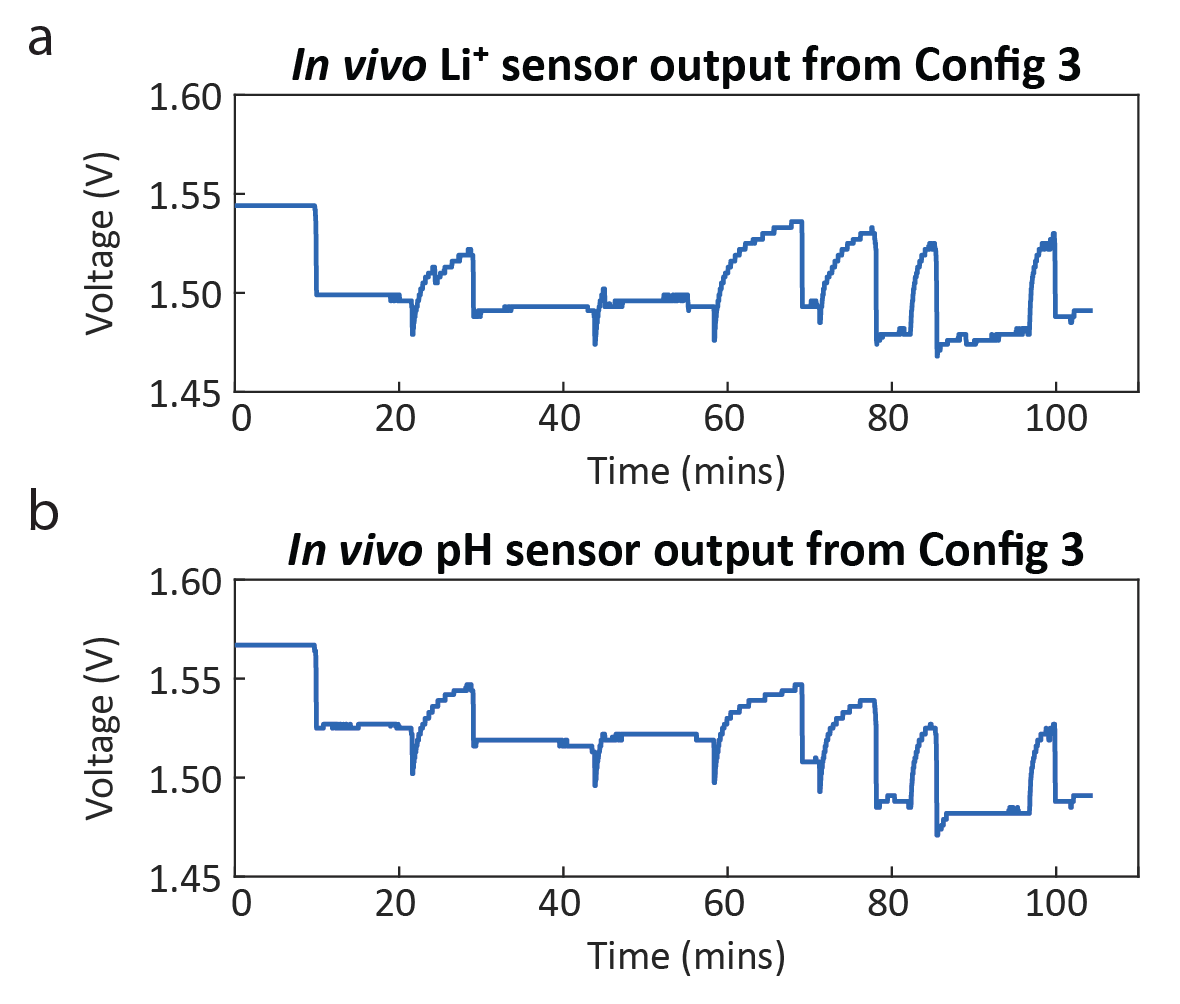

**S16.** **Sensor data from Config 3 of MiFi during *in vivo* experiment**. Both signals are filtered using moving median filter to remove high frequency noise. (a) Uncorrected Li sensor output. (b) Uncorrected pH sensor output.
